# Deletion of Lats 1 and 2 in mouse pancreatic acinar cells induces pancreatic fibrosis and inflammation

**DOI:** 10.1101/522938

**Authors:** Ming Gao, Jun Liu, Michael Nipper, Francis E. Sharkey, Randy L. Johnson, Howard C. Crawford, Yidong Chen, Pei Wang

## Abstract

**Objective:** The Hippo signaling pathway is known for regulating proliferation, differentiation, organ size, and tumorigenesis. Large tumor suppressor kinase 1 and 2 (LATS1&2) are the core kinases of this pathway, whose functions in both the normal pancreas and pancreatic diseases are unclear. We studied the function of LATS1&2 specifically in pancreatic acinar cells of adult mice.

**Design:** We generated mice with adult pancreatic acinar cell–specific deletion of *Lats1&2* genes by using CreER/LoxP. Pancreata were analyzed by histological examination, immunostaining, western blot, and RNA-sequencing.

**Results:** Deletion of *Lats1&2* genes in adult pancreatic acinar cells resulted in rapid development of pancreatic inflammation and fibrosis. Loss of Lats1&2 did not directly induce acinar cell proliferation or apoptosis, but resulted in pancreatic stellate cell (PSC) activation followed by immune cell infiltration and acinar-to-ductal metaplasia. These effects were mediated by the Hippo downstream effectors YAP1/TAZ. By using neutralizing antibody to block CTGF, a YAP1/TAZ target, the inflammation and fibrosis were reduced. Our RNA-sequencing data identified upregulation of fibroinflammatory genes in *Lats1&2* null pancreata, which may play important roles in stimulating PSC activation and promoting pancreatic fibrosis, as well as inflammation.

**Conclusions:** Deletion of the *Lats1&2* genes from adult acinar cells leads to the YAP1/TAZ dependent upregulation of a fibroinflammatory program. Our results emphasize the critical role of Lats1&2 in regulating PSC activation. Our findings identify new strategies for controlling pancreatic inflammation and fibrosis in diseases such as pancreatitis and pancreatic cancer.

## Introduction

Inflammation is a protective mechanism to clear damaged tissues and plays essential roles in tissue repair after injury. However, an uncontrolled inflammatory response can cause severe tissue injury[1]. Recent clinical data support the existence of continuum between recurrent acute pancreatitis (AP) and chronic pancreatitis (CP)[2]. CP is caused by a persistent inflammatory response that eventually leads to fibrosis and pancreatic failure. Currently, there are no specific treatments or prevention strategies for this disease[3]. While premature intra-acinar trypsinogen activation has been considered as a major pathogenic factor contributing to AP[4], the mechanisms driving pancreatic fibrosis, the characteristic feature of CP, during the progression from acute pancreatitis to chronic pancreatitis, remain unclear[5].

Extensive evidence suggests that activation of pancreatic stellate cells (PSCs) plays the central role in pancreatic fibrosis[6]. Although a number of studies have shown a passive role of damaged acinar cells in activating PSCs via recruitment of immune cells, the possibility that acinar cells may actively modulate the micro-environment to support sustained PSC activation has not been well studied[7],[8]. Indeed, several key PSC activators such as TGF-β and CTGF are upregulated in acinar cells adjacent to the areas of fibrosis in pancreata of chronic pancreatitis patients[9],[10]. Therefore, it is important to understand the mechanisms mediating acinar cell-PSC communication and to investigate their contribution to the pathogenesis of chronic pancreatitis.

As the central kinases of the Hippo signaling pathway in mammals, large tumor suppressor 1 and 2 (LATS1&2) play important roles in inhibiting proliferation, promoting apoptosis, and inhibiting F-actin polymerization[11]. LATS1&2 directly phosphorylate downstream effectors YAP1/TAZ and restrict the activity of YAP1/TAZ via both cytoplasmic retention and ubiquitin-proteasome-mediated degradation. Once LATS1&2 is inactivated, unphosphorylated YAP1/TAZ translocate to the nucleus to initiate the transcription of many genes, including pro-proliferative and anti-apoptotic genes[12]. However, recent studies suggest that the functions of the Hippo pathway are highly context dependent. In sharp contrast to the overgrowth phenotype in mice with hepatic Hippo inactivation, inactivation of the Hippo pathway in the developing pancreas leads to a decrease of pancreatic mass due to postnatal dedifferentiation of acinar cells[13],[14]. Nevertheless, in the cellular context of the adult pancreas, the roles of Hippo pathway in pancreatic homeostasis are unknown. The fact that the expression of YAP1 and TAZ has been found in human pancreatic cancer and pancreatitis samples[15],[16] strongly suggests that the Hippo pathway may play important roles in the pathogenesis of pancreatic disease.

In this study, we confirmed that the core components of the Hippo pathway dynamically changed in experimental pancreatitis models. By taking advantage of the Ptf1a^Cre-ER^/LoxP system in mouse pancreas, we found that mice with acinar-specific deletion of Lats1&2 developed strong inflammation and surprisingly rapid and severe fibrosis in a YAP1/TAZ dependent manner. We further identified a number of upregulated pro-inflammatory and pro-fibrotic genes in Lats1&2 deficient cells, suggesting that these identified genes might be targets of YAP1/TAZ. Our data also suggest that the Hippo signaling pathway can mediate acinar cell-PSC communication during the pathogenesis of pancreatitis.

## Materials and Methods

### Generation of conditional knockout mice

All animal study protocols were approved by the UT Health San Antonio Animal Care and Use committee. *Ptf1a^Cre-ERTM^* mice (stock number: 019378) and *R26R-EYFP* mice (stock number: 006148) were obtained from Hebrok Lab[17]. *Lats1^fl/fl^* and *Lats2^fl/fl^* mice were kindly provided by Dr. Randy L. Johnson. *Yap1^fl/fl^* and *Taz^fl/fl^* mice were kindly provided by Dr. Eric N. Olson. We generated (1) *Ptf1a^CRE-ER^Rosa26^LSL-YFP^* mice (P mice, control), (2) *Ptf1a^CRE-ER^Rosa26^LSL-YFP^Lats1^fl/fl^Lats2^fl/fl^* mice (PL mice), (3) *Ptf1a^CRE-ER^Yap1^fl/fl^Taz^fl/fl^* mice (PTY mice), and (4) *Pff1a^CRE-ER^Rosa26^LSL-YFP^Lats1^fl/f^lLats2^fl/fl^Yap1^fl/fl^Taz^fl/fl^* mice (PLTY mice).

### CTGF inhibition experiment

PL mice were injected once with neutralizing mAb FG-3019 (30mg/kg, FibroGen) by intraperitoneal (i.p.) administration. Twenty-four hours later, mice were subjected to 180mg/kg of TAM once, and then continued to be treated with FG-3019 every other day. Human IgG (equal amount, Jackson ImmunoResearch) was used as the control. Mice were sacrificed at Day 15 after TAM injection. Pancreatic tissue sections were stained with H&E to quantify the severity of inflammation.

### Statistical analysis

All data are expressed as mean ± standard error of the mean. GraphPad Prism 5 Software was used to analyze the data with ANOVA and Student’s t-test.

### Additional Methods

Detailed methods can be found in the Supplementary Materials and Methods section.

## Results

### The Hippo pathway is dynamically regulated during caerulein-induced pancreatitis mouse models

To test potential involvement of the Hippo pathway in pancreatitis, we examined the histological alterations and the expression patterns of Hippo pathway core components in time course analysis of caerulein-induced AP (Figure 1A). Pancreatic inflammation was observed from Day 1 to 3, and recovery started at Day 4. By Day 6, the pancreas appeared normal (Figure 1B). Interestingly, western blot results revealed that the protein levels of Lats1&2 were significantly downregulated from Day 1. By Day 6, the expression of both LATS1 and LATS2 recovered. In addition, the level of phospho-YAP1 (P-YAP1) showed a similar trend as LATS kinase. Inversely, YAP1/TAZ were increased from Day 1, and when the pancreas recovered, YAP1/TAZ went down back to normal levels (Figure 1C, Supplementary Figure 1A). YAP nuclei staining was observed at Day 1 in the AP pancreas (Supplementary Figure 1B). These data indicated a strong relationship between the Hippo signaling pathway and inflammatory response. CP was induced by repetitive episodes of AP for four weeks (Figure 1D), resulting in pancreatic fibrosis and increased collagen deposition, which was stained by Sirius Red (Figure 1E). Western blot data clearly showed significant reduction of LATS1&2 levels and elevated levels of YAP1/TAZ in the CP group when compared to control mice (Figure 1F, Supplementary Figure 1C), suggesting that Hippo signaling is also involved in pancreatic fibrosis. Taken together, we found that the Hippo signaling pathway is dynamically regulated during caerulein-induced pancreatitis, indicating a potential function of Hippo signaling in modulating pancreatic inflammation and fibrosis.

**Figure 1.**
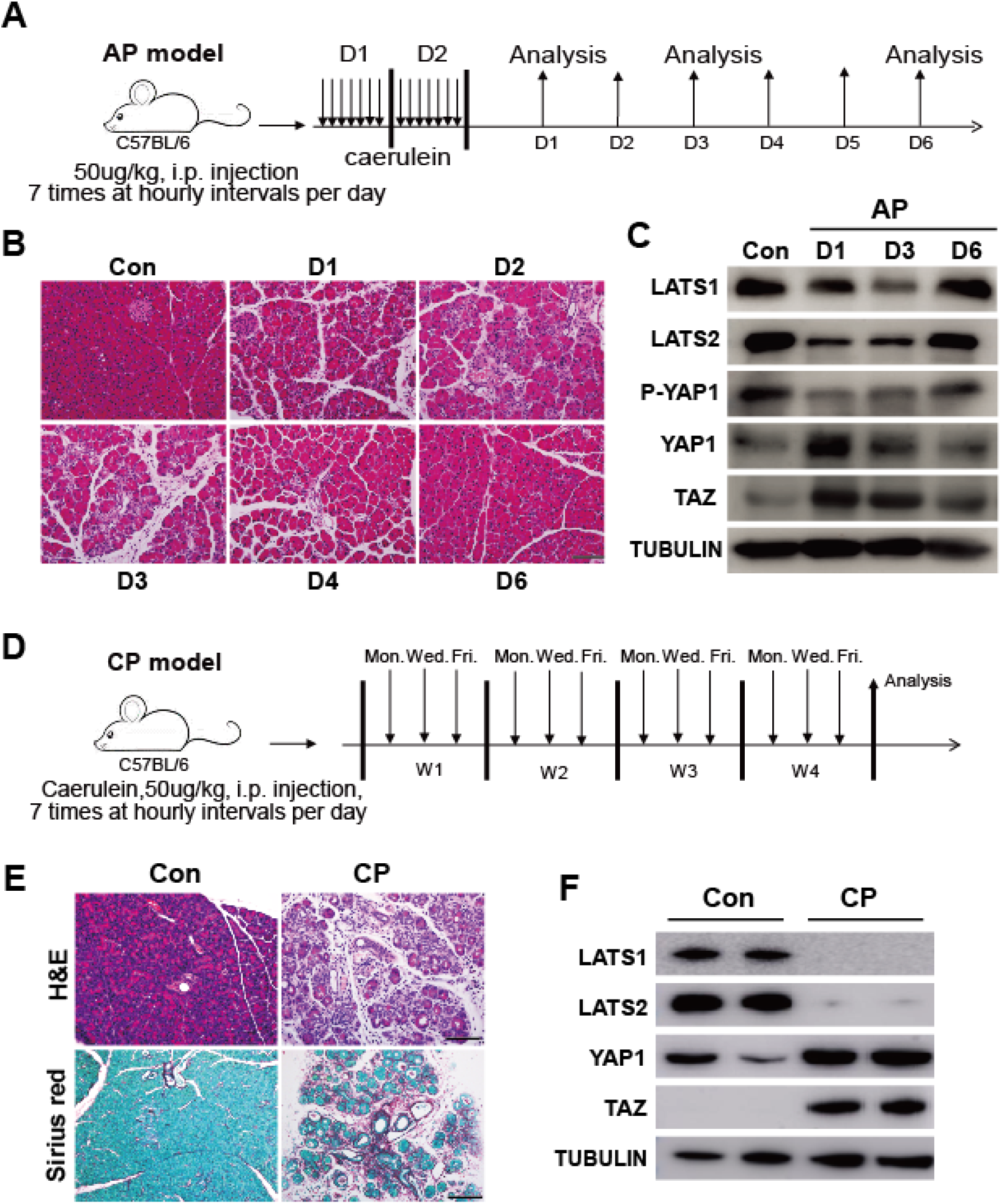
The Hippo signaling pathway is involved in inflammation and fibrosis in Caerulin-induced pancreatitis mouse model. (A) Schematic diagram of induction of acute pancreatitis (AP). n=3. (B) The acute pancreatitis was analyzed by H&E examination on D1, D2, D3, D4, and D6 after the induction. (C) Western blot of LATS1, LATS2, P-YAP1, YAP1, and TAZ in untreated and AP mice. Untreated mice served as the control group. Tubulin was used as the internal control. (D) Schematic diagram of induction of chronic pancreatitis (CP) (n=4). (E) H&E examination of CP mice. Untreated mice served as the control group. Sirius Red was used to detect collagen deposition in CP mice. (F) Western blot of LATS1, LATS2, YAP1, and TAZ in untreated and CP mice.

### Loss of LATS1&2 in pancreatic acinar cells causes pancreatic inflammation and fibrosis in adult mice

To investigate the roles of LATS1&2 in the pancreas, we genetically ablated *Lats1* and *2* genes specifically in pancreatic acinar cells at the adult stage. Through multiple steps of breeding (Supplementary Figure 2A), we generated *Lats1* and *2* double knockout mice (PL: *Pf1a^CreER^Lats1^fl/fl^Lats2^fl/fl^Rosa26^LSL-YFP^*) and *Lats1* and *Lats2* single knockout mice (PL1KO: *Ptf1a^CreER^Lats1^m^Lats2^fl/+^Rosa26^LSL-YFP^;* PL2KO: *Ptf1a^CreER^Lats1^fl/+^Lats2^fl/fl^Rosa26^LSL-YFP^*). We used both littermate (L1/2: *Ptf1a^+/+^Lats1^fl/fl^Lats2^fl/fl^Rosa26^LSL-YFP^*) and Cre alone (P: *Ptf1a^CreER^Rosa26^LSL-YFP^*) mice as controls. We administered 180mg/kg/day of tamoxifen (TAM) to 6-12 week old mice via intraperitoneal (i.p.) injection for five consecutive days. We found that PL mice become weaker and start losing weight by seven days after the last TAM injection (Supplementary Figure 2B). At day 10, PL mice had a weight loss of 19.2%. We found a significant reduction of blood glucose levels in PL mice, suggesting a starvation phenotype (Supplementary Figure 2C). We observed a continuous loss of acinar tissue and replacement by fibrous tissue over 14 days (Supplementary Figure 2D).

Histologically, PL mice exhibited severe damage in the pancreas, including acinar cell atrophy, inflammatory cell infiltration, and acinar-to-ductal metaplasia (ADM) at Day 5 after the last injection of TAM (Figure 2A, 2D, 2E). No such defects were observed in the pancreata of control groups (Figure 2A). Successful *Lats1&2* deletions in the pancreas were determined by PCR analysis (Supplementary Figure 3A) and western blot (Figure 2B). The protein levels of downstream effectors YAP1/TAZ were increased dramatically in PL mice, while phospho-YAP1 levels were lower in PL mice (Figure 2B). YAP1 and TAZ nuclear translocations were detected in PL mice by immunohistochemistry (IHC) staining (Figure 2C, Supplementary Figure 3B), suggesting that loss of LATS1&2 in pancreatic acinar cells induced YAP1/TAZ activation, similarly to other tissues [18]. Compared to the control mice, we detected an increased expression of ductal cell marker cytokeratin 19 (CK19) in PL mice (Figure 2D). YFP+/CK19+ cells were only observed in PL mice, suggesting that acinar cells underwent ADM, a common hallmark of pancreatitis (Figure 2D, Supplementary Figure 4A). Ki67 expression was found in YFP+/CK19+ cells, suggesting that proliferation was activated during ADM (Supplementary Figure 4B). We found apoptotic cells in PL mice by staining of cleaved caspase-3 (Supplementary Figure 4C). Consistent with H&E staining, we identified infiltration of immune cells in PL mice by CD45 staining (Figure 2E, Supplementary Figure 4A). We detected strong staining of α-smooth muscle actin (αSMA) and significant deposition of collagen in PL pancreata (Figure 2F, Supplementary Figure 4A), suggesting the occurrence of pancreatic stellate cell (PSC) activation and pancreatic fibrosis.

**Figure 2.**
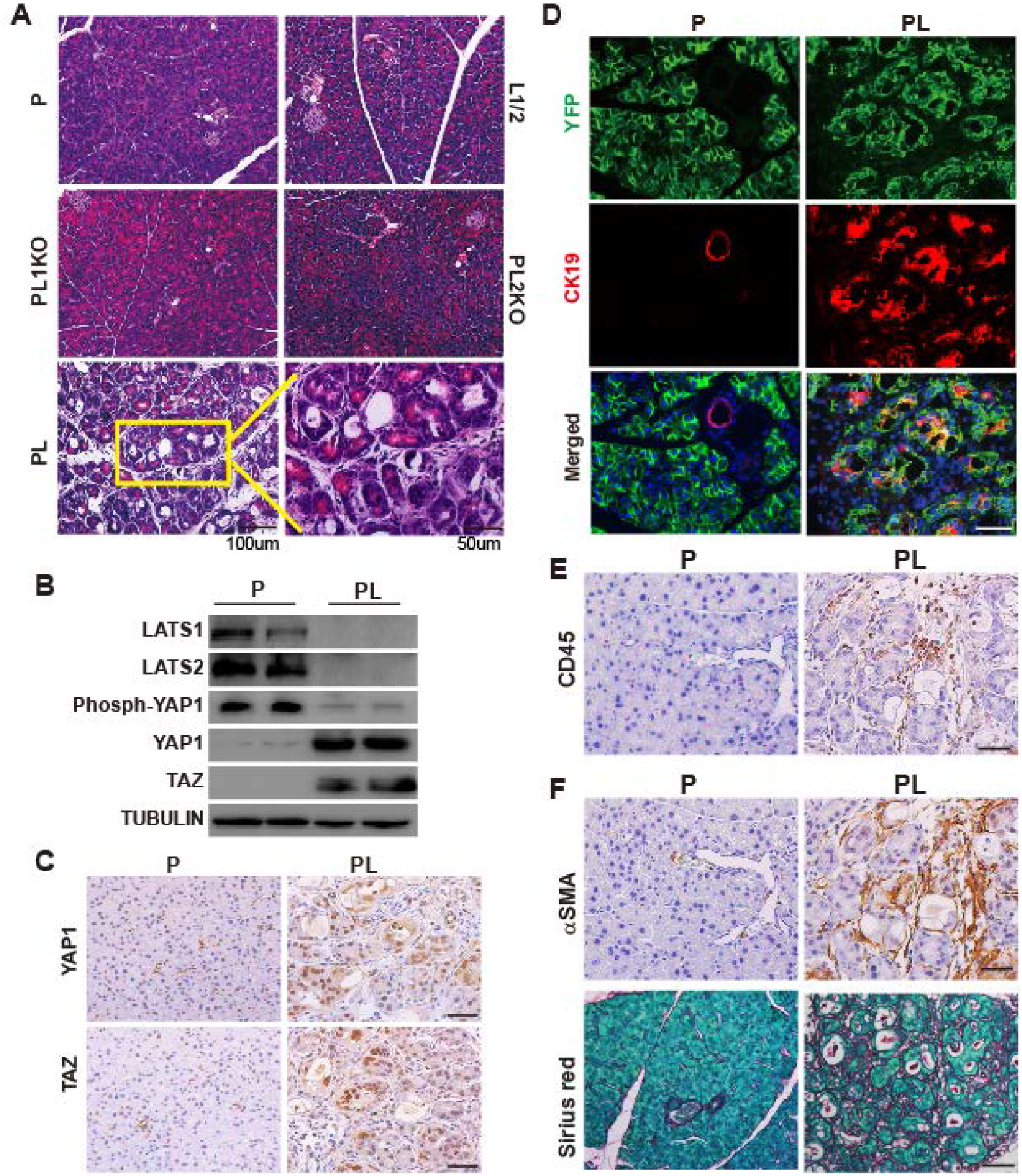
Acinar cell-specific Lats1&2 deletions caused pancreatic inflammation and fibrosis in adult mice. P, PL, L1/2, PL1KO, and PL2KO mice were administrated with 180mg/kg/day of TAM for five consecutive days via i.p., respectively (n=6). Pancreata were collected at Day 5 after the final injection. (A) Histological morphology of P, L1/2, PL, PL1KO, and PL2KO mice. L1/2, PL1KO, and PL2KO mice were littermate controls. (B) Western blot showing the levels of LATS1, LATS2, phospho-YAP1, YAP1, and TAZ in control (P) and double knockout (PL) mice. TUBULIN was used as the internal control. (C) YAP1 and TAZ IHC staining in P and PL mice. (D) Immunofluorescence of YFP (Green) and CK19 (Red) in P and PL mice. Nuclei stained with DAPI (Blue). (E) P and PL pancreata stained with anti-CD45 antibody. (F) P and PL pancreata stained with anti-*α*SMA antibody, and Sirius Red, respectively.

Taken together, these results strongly suggest that the loss of LATS1&2 in pancreatic acinar cells rapidly leads to pancreatic fibrosis, a major clinical feature of chronic pancreatitis.

### Loss of *Lats1&2* in pancreatic acinar cells stimulates PSC activation

Inflammation in the pancreas is frequently induced by acinar cell injury[19]. To test whether inflammation in PL mice was induced by autonomous acinar cell death following Lats1&2 deletions, we decided to knockout Lats1&2 in a small portion of acinar cells in the adult pancreas. If the inflammation was induced by acinar cell death due to Lats1&2 knockout, injured Lats1&2 null acinar cells would be rapidly removed by immune cells and the pancreas should recover with the remaining wild type acinar cells. We gave the mice single injection of different doses of TAM. *Ptf1a^CreER^* induced chromosome recombination in a TAM dose-dependent manner (see supplemental data and Supplementary Figure 5A, 5B). All mice developed inflammation 3 weeks after TAM injection and the severity of the inflammation was correlated with TAM dose (Supplementary Figure 5C). Strikingly, unlike in cerulein induced AP mice, the pancreatic lesions developed in low dose TAM injected PL mice did not recover over time. This long-lasting phenotype suggested that acinar damage was not the major mechanism to initiate inflammation in PL mice. The low dose (45mg/kg) TAM injection allowed us to examine acinar cell fate at the individual cell level. No increased apoptosis, proliferation, or ADM was observed in PL mice at 4, 8 or 12 days after TAM injection (see supplemental data and Supplementary Figure 6A-C), further supporting that loss of Lats1&2 did not directly induce autonomous acinar cell death.

To achieve a longer and clearer time window to analyze the temporal sequences of alterations in PL mice, we gave the mice a single TAM injection (180mg/kg) and analyzed the pancreata at Days 0, 5, 10, 15, and 20. The pancreata of PL mice almost disappeared by Day 20. At Day 15, the phenotype was similar to what we observed at Day 5 for mice with five injections of TAM (Figure 3A). CK19, αSMA, and CD45 staining by IHC in consecutive sections of PL mice and was positive in all lesions at Day 15, indicating the presence of ADM, PSC activation, and immune cell infiltration (Figure 3B, Supplementary Figure 7A). However, the small dispersed lesions at Day 10 in PL pancreata with YAP1/TAZ nuclear accumulation were surrounded by αSMA positive stromal cells, implying that PSCs might be activated by the direct communication with Lats1&2 null acinar cells (Figure 3C). Indeed, many of these lesions were not accompanied by CD45 positive cells (Figure 3D). This was further confirmed by co-staining αSMA with CD45 (Supplementary Figure 7B), highlighting the fact that PSCs were activated before significant infiltration of immune cells. These lesions retained acinar morphology and were negative for CK19, suggesting that PSCs were activated in the absence of ADM (Figure 3E). We isolated acinar cells from PL mice that received 5 injections of TAM and cultured them in a collagen-based 3D model. ADM was quantified by counting the number of newly formed duct-like structures in 3D culture[20],[21]. Lats1&2 deletions did not significantly enhance the efficiency of ADM in 3D culture without TGFα, but the size of ADM events increased in the presence of TGFα (Supplementary Figure 7C), suggesting that acinar cells with Lats1&2 deletions might be primed for ADM but require additional environmental signals to undergo ADM *in vivo*. Together, our data indicate that activation of PSCs was an early event in PL mice.

**Figure 3.**
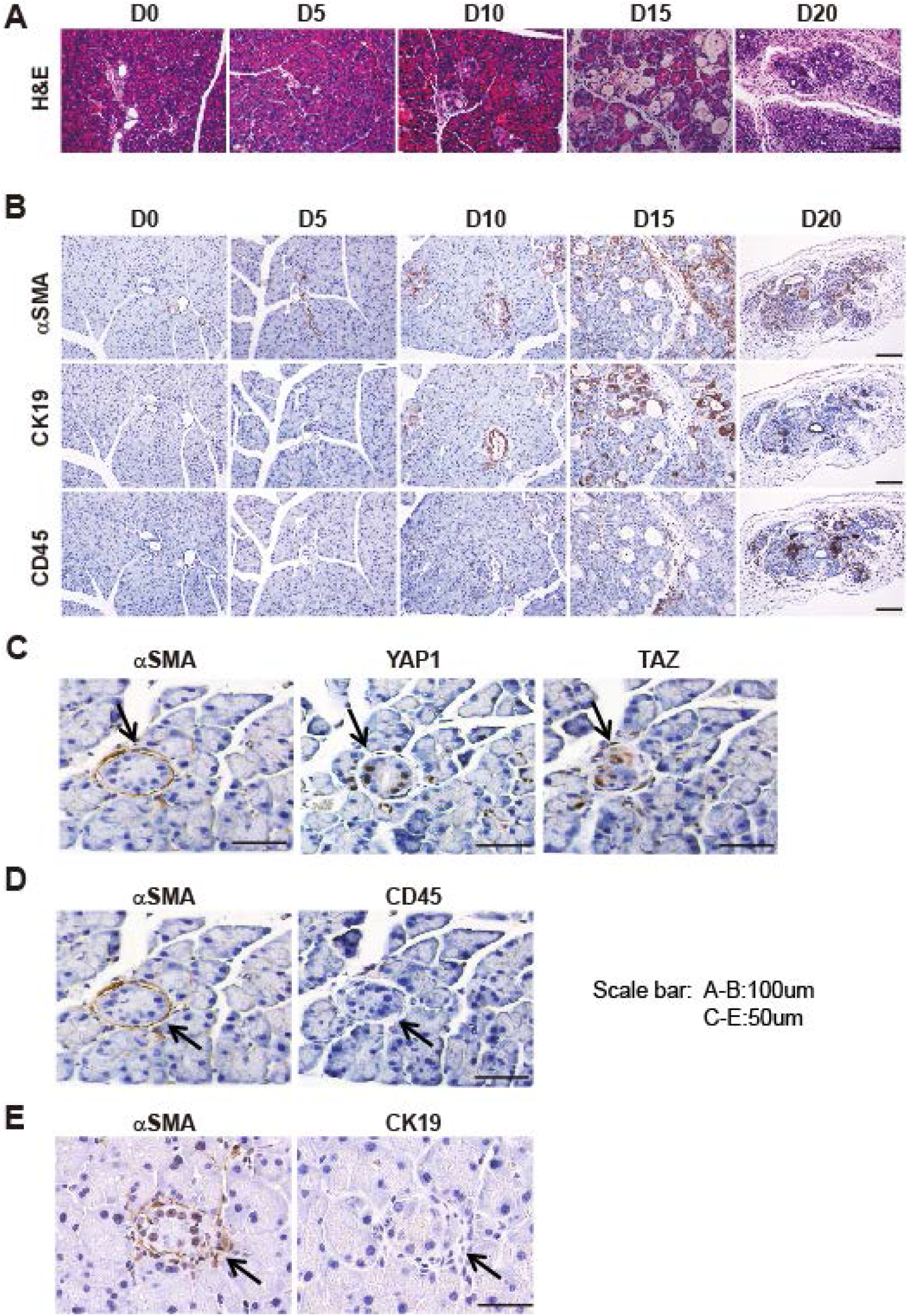
*Lats1&2* deletion in acinar cells results in PSC activation. PL mice were injected once with 180mg/kg of TAM, pancreata were collected and embedded at Day 0, Day 5, Day 10, Day 15, and Day 20 after injection, respectively (n=4-6). (A) Morphological changes over time (D0, D5, D10, D15, D20) were examined by H&E staining. (B) Anti-αSMA, anti-CD45, and anti-CK19 antibodies were used to stain activated PSCs, immune cells, and ductal-like cells, respectively. (C) αSMA, YAP1, and TAZ IHC staining in consecutive sections at Day 10. (D) αSMA and CD45 IHC staining in consecutive sections at Day 10. (E). αSMA and CK19 IHC staining in consecutive sections at Day 10

### *Lat1&2* deletion-induced YAP1/TAZ-dependent pancreatic inflammation and fibrosis

*Lats1&2* deletion in adult acinar cells dramatically upregulated YAP1/TAZ in the pancreas and induced YAP1/TAZ nuclear translocation (Figure 2B, 2C, Supplementary Figure 3B). Thus, we tested whether the inflammation and fibrosis in *Lats1&2* null pancreata were mediated by YAP1/TAZ. Loss of *YAP1/TAZ* in adult acinar cells *(Ptf1a^CreER^YAP1^fl/fl^TAZ^fl/fl^*, PTY) did not result in overt defect (Supplementary Figure 8A). We then crossed PL mice with *Yap1^fl/fl^Taz^fl/fl^* mice to generate *Ptf1a^CreER^Rosa26^LSL-YFP^Lats1^fl/fl^Lats2^fl/fl^Yap1^fl/fl^Taz^fl/fl^* (PLTY) mice. The PLTY mice were administered with 180 mg/kg/day of TAM for 5 consecutive days to induce the quadruple deletions of *Lats1&2* and *YAP1/TAZ* genes in adult pancreatic acinar cells. P and PL mice were used as controls. All pancreata were harvested at Day 5 after the final injection of TAM (Supplementary Figure 8B). The deletions of *Lats1, Lats2, Yap1*, and *Taz* were confirmed by western blot (Figure 4A, Supplementary Figure 8C). H&E staining revealed that ablation of *YAP1/TAZ* rescued the pancreatitis phenotype of PL mice (Figure 4B). PSC activation, immune cell infiltration, and ADM were remarkably reduced in PLTY mice when compared with PL mice (Figure 4C, Supplementary Figure 8D). These data strongly suggested that *Lats1&2* deletion induced inflammation and fibrosis depend on YAP1/TAZ.

**Figure 4.**
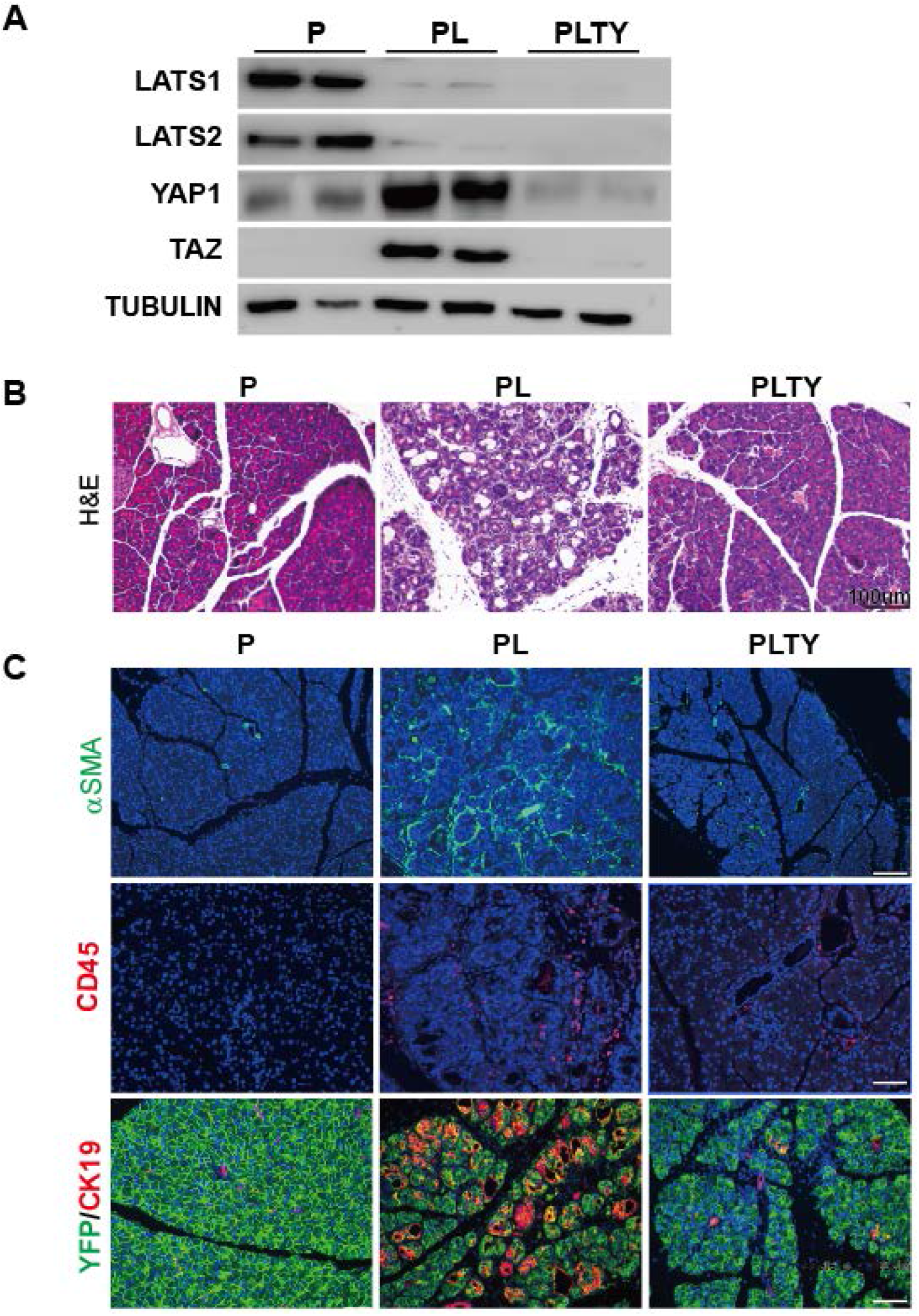
Loss of YAP1/TAZ in acinar cells rescues Lats1&2 deletion-induced pancreatic inflammation and fibrosis. P, PL, and PLTY mice were subjected to 180mg/kg/day of TAM for 5 consecutive days via i.p. (n=6). Pancreata were collected 5 days later. (A) Western blot of Lats1, Lats2, YAP1, and TAZ in P, PL, and PLTY mice. Tubulin was used as the internal control. (B) Histological examination by H&E staining among P, PL, and PLTY groups. (C) Immunofluorescence staining of αSMA (Green), CD45 (Red), and ADM (Green + Red) among P, PL, and PLTY groups. Nuclei stained with DAPI (blue).

### CTGF inhibition attenuates *Lats1&2* deletion-induced pancreatitis

After genetically confirming that the pancreatitis phenotype caused by loss of *Lats1&2* is mediated through YAP1/TAZ activation, we investigated how YAP1/TAZ activation triggered pancreatitis. Connective tissue growth factor (CTGF) is the best known YAP1/TAZ target gene and has been documented to play a central role in pancreatic stellate cell activation[22] and fibrosis[23]. We found that *Ctgf* mRNA expression was up-regulated at an early time point (D2) after *Lats1&2* deletions (Figure 5A). In addition, CTGF protein level was significantly decreased in PLTY mice compared to PL mice (Figure 5B, Supplementary Figure 9A). To determine whether CTGF was responsible for pancreatic fibrosis in PL mice, we performed a rescue experiment using CTGF-neutralizing antibody FG-3019 (FibroGen, Inc.)[24]. PL mice were injected with FG-3019 or FG-hulgG (control) for 2 days (ip. 30mg/kg/day), followed by a single TAM (180mg/kg) injection. After this, the mice received FG-3019 or FG-hulgG injection every other day and were sacrificed at Day 15 (Figure 5C). The H&E staining revealed that CTGF depletion partially attenuated inflammation and fibrosis in PL mice (Figure 5D). FG-3019 treated mice achieved 61.7%, 53.5% and 56.7% reduction of immune cell infiltration, stromal reaction and ADM, respectively, compared with the FG-hulgG treated group (Supplementary Figure 9B), suggesting that neutralization of CTGF partially rescued the pancreatitis induced by *Lats1/2* deletions in adult acinar cells. To examine whether neutralizing CTGF affected the health of the mice, we treated another group of mice (n=6) for 30 days. Half of FG-3019 treated mice maintained their body weight, while all FG-hulgG treated mice lost weight by day twelve (Supplementary Figure 9C).

**Figure 5.**
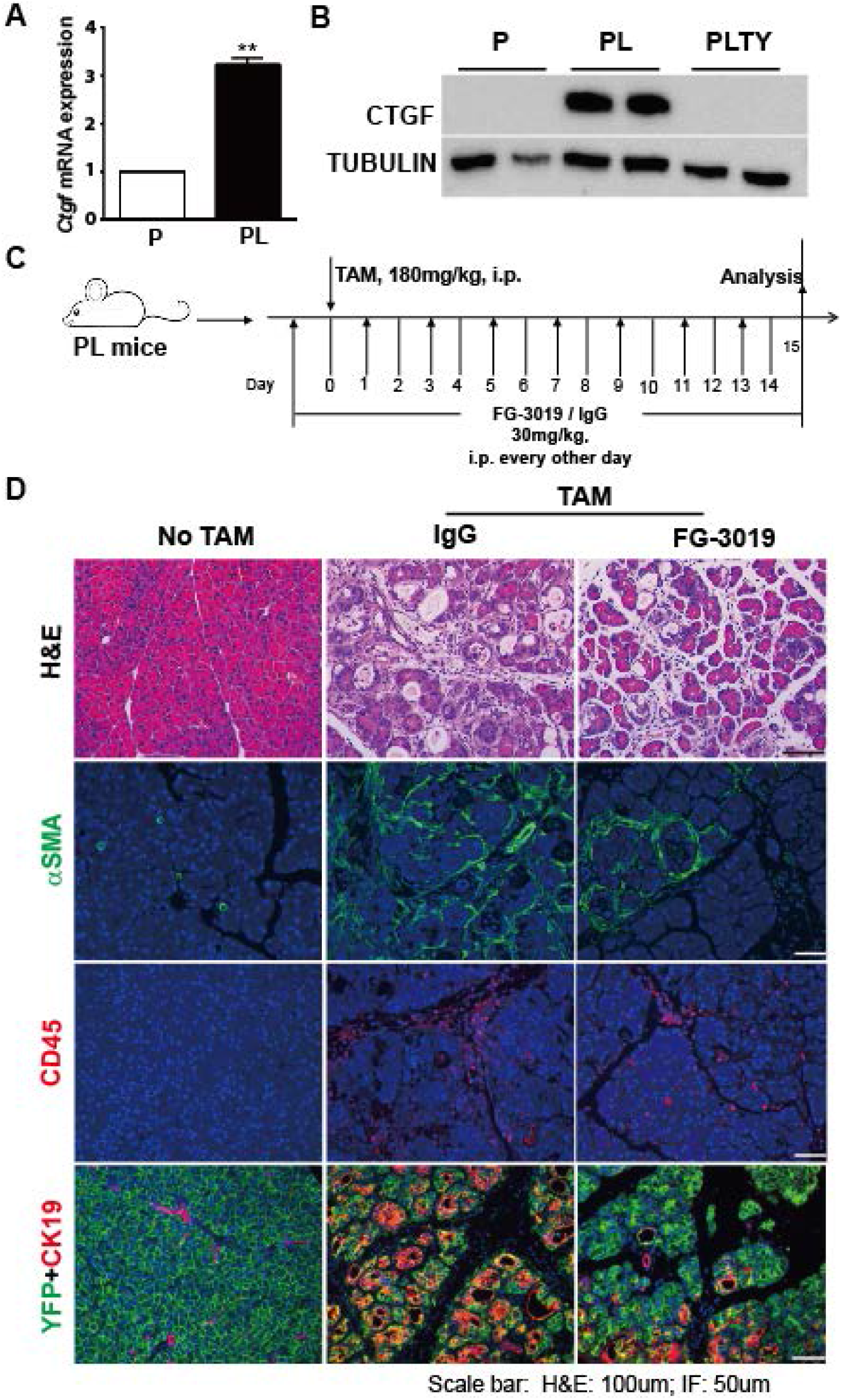
Depletion of CTGF partially abolishes pancreatic inflammation and fibrosis in *Lats1&2* double knockout mice. (A) *Ctgf* mRNA expression in P and PL mice (n=3). **, P<0.01. (B) Western blot of CTGF protein levels in P, PL, and PLTY mice. TUBULIN served as the internal control. (C) Schematic diagram of experimental design (n=6 per group). Mice without TAM injection served as control group. FG-3019: CTGF neutralizing antibody; IgG: human control IgG. (D) Top: Histological examination by H&E staining; Center: Immunofluorescent staining of αSMA (Green); Bottom: Immunofluorescent staining of CD45 (Red). Nuclei stained with DAPI (blue).

### Loss of *Lats1&2* induces upregulation of *pro-inflammatory and pro-fibrogenic* genes in the pancreas

The partial rescue achieved by CTGF depletion suggested that other targets of YAP1/TAZ may contribute to the development of inflammation and fibrosis in PL mice. Thus, we performed RNA sequencing analysis of pancreata from control P and mutant PL mice. To increase the number of *Lats1&2* null cells, control and PL mice were subjected to180mg/kg/day of TAM for five consecutive days. Tails of pancreata were harvested at Days 1, 2, and 3 (n=3-5 for each time point) for RNA extraction and RNA-seq library preparation. Histological examination showed no visible defects in Day 1, 2 or 3 PL pancreata (Supplementary Figure 10A). There was no positive staining of CD45, αSMA, or CK19 at Day 2, but there was some at Day 3 (Supplementary Figure 10B).

We compared transcriptomes from RNA-seq data from PL mice at Days 1, 2, and 3 after TAM injection to control mice and obtained 40, 405, and 765 differentially expressed genes, respectively, using the significance criterion (fold-change > 2; adjusted p-value < 0.05, and RPKM > 2). A large number of genes with expression changes were found in Day 2 PL pancreata (Figure 6A), although no significant PSC activation or immune cell infiltration were observed at this time point. We performed K-means clustering analysis and identified two representative clusters: 300 upregulated genes and 232 downregulated genes (Figure 6B). Several known YAP1/TAZ targets including *Ctgf, Cyr61*, and *Spp1* were among the upregulated genes (Supplementary Figure 10C). Gene Ontology GO term analysis with the 300 upregulated genes showed that five out of the top ten GO terms were related to immune response, which is consistent with the phenotype of PL mice. On the other hand, downregulated genes were related to ER and digestion, which are associated with normal acinar cell function (Figure 6B).

**Figure 6.**
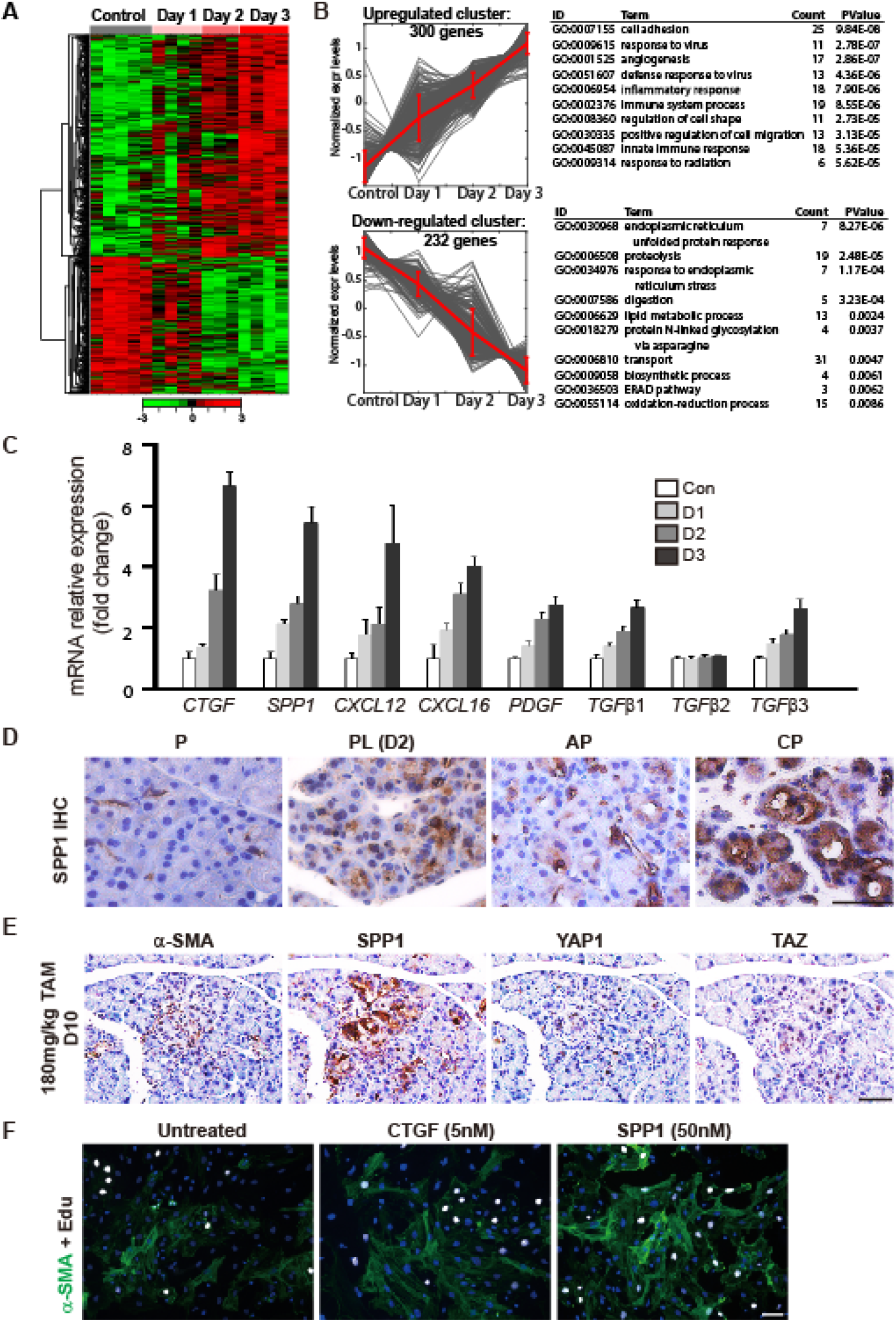
RNA-sequencing analysis of LATS1&2 regulated genes in mouse pancreas. (A) Heat map analysis showed differential gene expression patterns at different days (n=3-5). (B) Left: K-means clustering analysis showed the gene expression patterns of two classes (upregulated and downregulated genes) that were changed with time. Right: GO term analysis of the function of top upregulated and downregulated genes. (C) Validation of pro-fibrotic gene expressions by Q-PCR. (D) SPP1 IHC staining in P, PL (D2) mice, acute pancreatitis, and chronic pancreatitis mice. (E) The small lesions were stained with anti-αSMA, anti-SPP1, anti-YAP1, and anti-TAZ antibodies in consecutive sections. (F) Isolated PSCs were treated with CTGF (5nM) or SPP1 (50nM) for 3 days (n=3). Untreated group serves as the control. Anti-αSMA antibody was used to stain activated PSCs (Green). Cell proliferation was evaluated by Edu incorporation (White). Nuclei stained with DAPI (Blue).

We specifically analyzed GO terms regarding growth factor activity (GO:0008083), cytokine activity (GO:0005125), and chemokine activity (GO:0008009), and identified a total of 12 upregulated genes (*Bmp1, Ctgf, Cxcl12, Mia, Pdgfα, Tgfβ1, Tgfβ3, Timp1, Ccl12, Cxcl10, Cxcl14, Spp1*). Interestingly, several genes involved in PSC activation *(Tgfβ* and *Pdgfα)* and fibrogenesis *(Spp1, Ctgf, Cxcl12*, and *Cxcl16)* were significantly upregulated at Day 2, before immune cell infiltration, suggesting a potential function of acinar cell YAP1/TAZ in direct regulation of PSC activation and pancreatic fibrosis. We validated the expression changes of *Ctgf, Spp1, Cxcl12, Cxcl16, Pdgfα, TGFβ1, TGFβ2*, and *TGFβ3* by Q-PCR (Figure 6C). Except for *TGFβ2*, all of these genes showed gradual upregulation during the time course, strongly supporting that they may be direct targets of YAP1/TAZ.

Among these genes, the upregulation of *Spp1* (secreted phosphoprotein 1) was the most dramatic. Even at Day 1 after TAM injection, we detected an about two-fold increase of *Spp1* compared to the control. SPP1 is expressed in pancreatic ductal tissues and undifferentiated pancreatic precursors[25], as well as in pancreatic cancer[26]. However, its role in pancreatitis is unknown. We found SPP1 positive staining in ductal cells in control pancreata, but also in acinar cells in caerulein-induced AP and CP, with much stronger staining in CP pancreata (Figure 6D). *Spp1* has been suggested as a potential target of YAP1/TAZ[27], and is also strongly associated with TGFβ function and hepatic stellate cell (HSC) activation[28]. Thus, we performed immunostaining of SPP1 on consecutive sections of day 10 PL pancreata with one dose of TAM (180mg/kg). We found that SPP1 staining correlated well with αSMA, YAP1, and TAZ staining in small lesions (Figure 6E). SPP1 was strongly positive while no apparent αSMA staining surrounded acinar cells at day 2 after five doses of TAM (Supplementary Figure 10D). The SPP1 positive cells were obvious in small lesions in day 10 PL pancreata (Supplementary Figure 10E). Thus, *Spp1* is likely both a target of YAP1/TAZ in pancreatic acinar cells and involved in PSC activation through YAP1/TAZ activation. To further test whether SPP1 can activate PSCs, We isolated the primary PSCs by density gradient centrifugation to directly test the effect of SPP1 on PSC activation [29]. The purity of the PSCs was confirmed by GFAP staining (Supplementary Figure 11A). The primary PSCs were treated with SSP1 recombinant proteins (50nM) for 3 days. We used CTGF recombinant protein as the positive control. SPP1 treatment significantly increased α-SMA positive cell numbers, similarly to what we observed in the CTGF treated group, suggesting that SPP1 can trigger PSC activation (Figure 6F, Supplementary Figure 11B). SPP1 treatment did not significantly enhance the proliferation of PSCs as determined by Edu staining (Supplementary Figure 11B). We noticed that α-SMA positive cells and Edu positive cells did not always overlap (Figure 6F), suggesting that they might be two uncoupled events in PSC activation. Together, these data supported the notion that SPP1 might be an important soluble factor secreted by acinar cells with Lats1&2 deletions to activate PSCs.

### M2 macrophages are dominant in the *Lats1&2* null pancreas

Immune cell infiltration increased dramatically in day 4 PL pancreata after five dose of TAM injection (Supplementary Figure 12A). Unlike the tissue damage-induced M1 polarization of macrophages in AP, alternatively activated macrophages (M2) are dominant in CP and activated PSCs are important inducers of M2 polarization[30]. Thus, we examined which macrophages were present in PL mouse. F4/80 staining revealed an increased number of macrophages in the pancreata of PL mice (Figure 7A). We used flow cytometry to sort CD45+CD11b+F4/80+ macrophages from the pancreata of P and PL mice four days after final TAM injection to examine the expression of a panel of M1- and M2-accociated genes (Supplementary Figure 12B). Compared with the controls, macrophages from PL mice displayed a dramatic upregulation of M2 macrophage-associated gene *Ym1* (Figure 7B). Other M2 macrophage-associated cytokines and chemokines including *Il10, Tgf-β1*, and *Ccl17* also showed significant up-regulation (Figure 7B). M1 macrophage-associated genes such as *Tnf-α, Sosc1*, and *Nos2* showed no significant expression change (Figure 7C) and *Il-1β* was barely detectable by Q-PCR. These data revealed that M2 macrophage polarization was rapidly induced upon YAP1/TAZ activation in adult acinar cells.

**Figure 7.**
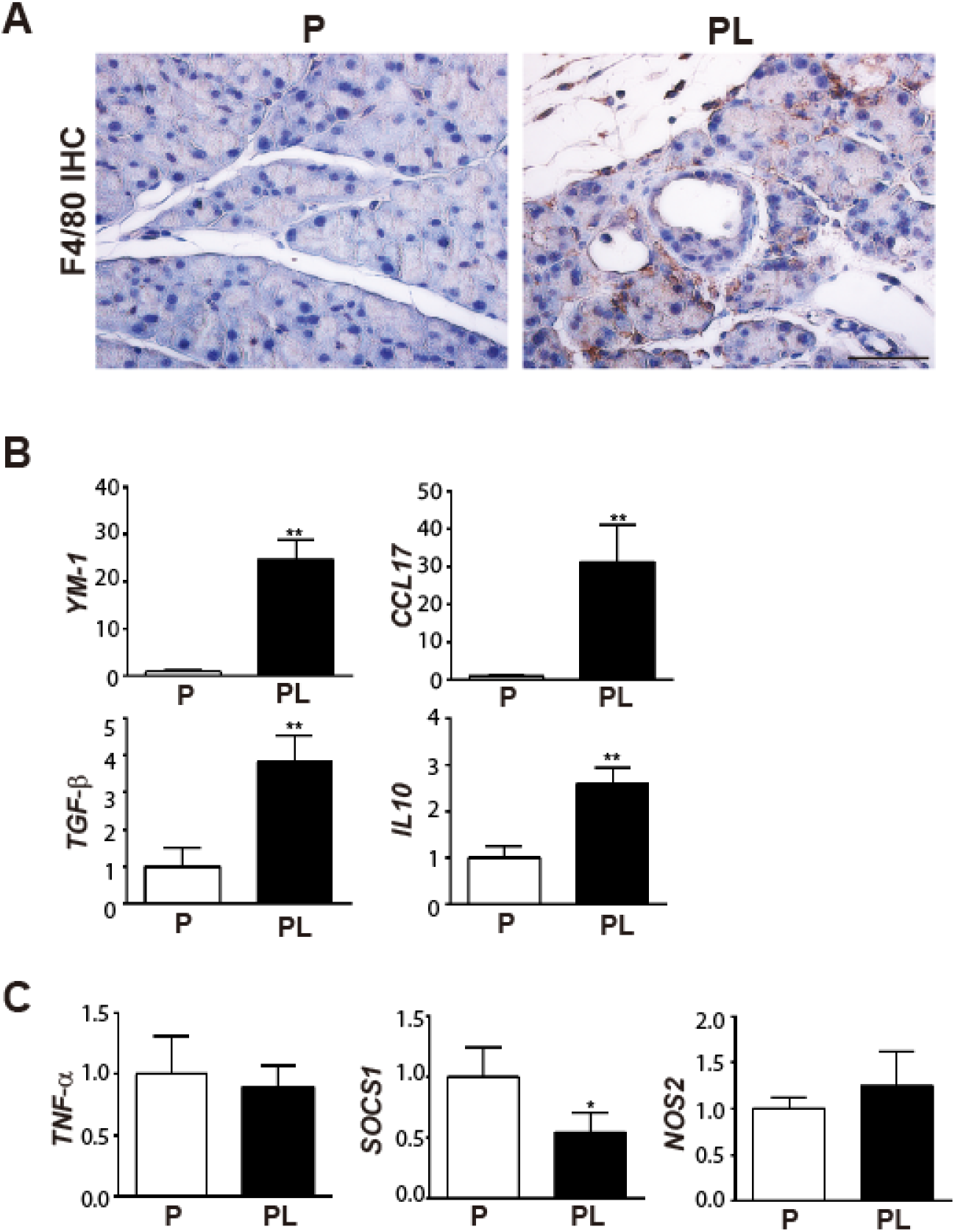
Loss of *Lats1&2* induces M2 macrophage polarization. P and PL mice were treated with 180mg/kg/day of TAM for 5 consecutive days, cells were isolated from both groups 4 days later (n=3). (A) F4/80 IHC staining in P and PL mice on Day 4 after TAM injection. (B) M2 macrophage specific genes *Ym-1*, *Ccl17*, *Tgf-β*, and *IL10* were elevated in sorted macrophages of PL mice. (C) M1 macrophage specific genes *Tnf-α*, *Socs1*, and *Nos2* were not significantly changed. **, P<0.01, *, P<0.05

## Discussion

In this study, we generated a mouse model with deletion of Hippo core kinases LATS1&2 in adult pancreatic acinar cells. These mice developed severe pancreatic fibrosis and inflammation at a strikingly fast speed which has not been seen in other mouse models[31],[13],[14]. This unique phenotype is mediated by the downstream effectors YAP1/TAZ, indicating that pro-inflammatory and pro-fibrotic genes might be targets of YAP1/TAZ. Our study revealed a novel player in the pathogenesis of chronic pancreatitis.

As a digestive enzyme production factory, when acinar cells are damaged, they may release enzymes which can cause severe detrimental effects in the abdominal cavity. It is plausible to speculate that acinar cells should have an intrinsic mechanism to quickly sense these changes and modulate their microenvironment to contain the damage and promote regeneration. Increasing numbers of studies have begun to investigate the active roles of acinar cells in regulating the inflammatory response[8],[32],[33]. Considering the fact that the Hippo pathway is regulated by soluble factors, cell-cell contact, and extracellular matrix (ECM) stiffness[34], it is tempting to speculate that acinar cells may utilize this pathway to sense these exogenous stimuli and actively translate them into signals to orchestrate the surrounding microenvironment. Indeed, we observed that YAP1/TAZ activated acinar cells are surrounded by activated PSCs at early time points of acinar-specific Lats1&2 deletion. These data highlight the possibility that the Hippo pathway plays a critical role in mediating the direct interactions between acinar cells and stromal cells to modulate the pancreatic environment. The roles of viable acinar cells in the direct modulation of the inflammatory environment have also been reported by Cobo et al.[35]. They showed that NR5A2, a key transcriptional factor previously known to maintain acinar identity, controls an inflammatory program in acinar cells. Nevertheless, our RNA-seq data showed that *Lats1&2* knockout did not affect the expression of *Nr5a2* in acinar cells, suggesting that *Nr5a2* downregulation was not responsible for the inflammation initiation in our acinar specific *Lats1&2* knockout model.

A recent study discovered that *Lats1&2* knockout cancer cells secreted DNA containing exosomes to promote Th1 response and created an anti-tumor inflammatory environment[36]. However, we found rapid accumulation of Th2 response-associated M2 macrophages in our mouse model, suggesting a different mechanism. Acinar cells have been thought to have a passive role in promoting inflammation via release of damage-associated molecular patterns when damaged[37]. In this sense, it is unexpected that deletion of *Lats1&2* in adult acinar cells would induce sterile inflammation, because it is not intuitive to connect tissue damage to the knockout of *Lats1&2* based on their well-known functions[38]’. Indeed, time course analysis revealed that PSC activation and immune cell infiltration happened well before acinar cell death after *Lats1&2* deletion. In addition, unlike the rapid tissue recovery from AP caused by acinar cell damage, deleting Lats1&2 in a small fraction of adult acinar cells by low dose TAM administration induced persistent pancreatic inflammation, suggesting that these cells were not rapidly removed by immune cells. These data strongly suggest that inflammation and fibrosis in acinar specific *Lats1&2* deletion pancreata are not initiated by acinar cell damage. Nevertheless, whether the acinar cells with *Lats1&2* deletions are more susceptible to apoptosis under an inflammatory microenvironment should be further examined, as this might help to explain the severe phenotype in our PL model.

We confirmed that the pancreatic fibrosis and inflammation in *Lats1&2* null pancreata is dependent on YAP1/TAZ. PSC activation occurs before immune cell infiltration, suggesting that PSCs can be directly activated by *Lats1&2* null adult acinar cells to promote fibrosis. Dysregulated PSC activation results in excessive deposition of ECM in the gland, contributing to the progressive fibrosis associated with chronic pancreatitis and pancreatic cancer[39]. Therefore, understanding the mechanisms regulating PSC activation offers potential therapeutic targets for the prevention and treatment of these diseases.

The ability of *Lats1&2* null acinar cells to directly activate PSCs provides an explanation for the rapid progression of pancreatic fibrosis in our model. Acute pancreatic damage induces the infiltration of M1 macrophages and PSCs are transiently activated. A recent study showed that activated PSCs polarize infiltrated M1 macrophages towards M2, which in turn can more efficiently activate the PSCs[30]. This “feed-forward” process between PSCs and macrophages promotes chronic pancreatitis-associated fibrosis. In PL pancreata, the persistent PSC activation supported by *Lats1&2* null acinar cells accelerates macrophage M2 polarization and rapidly creates a micro-environment in favor of fibrosis development. Our mouse model mimics the persistent Hippo pathway inactivation in chronic pancreatitis. Thus, acinar cell-mediated PSC activation might be one underlying mechanism contributing to the sustained fibrosis and inflammation in chronic pancreatitis even after the clearance of damaged tissue.

Of note, the concept that YAP1/TAZ-mediated epithelia-myofibroblast interaction plays an important role in tissue fibrosis may be applied to other tissues. TAZ induces the expression of Indian hedgehog (Ihh) to activate HSCs and promote nonalcoholic steatohepatitis-associated fibrosis and inflammation[40]. However, Ihh does not activate the proliferation or ECM synthesis of PSCs[41]. Therefore, *Lats1&2* null adult acinar cells may utilize other signaling molecules to activate PSCs. We found that the PSC activator CTGF, a well-known YAP1/TAZ target, was significantly upregulated in *PL* mice, corroborating with the clinical observation that CTGF expression is induced in acinar cells in human chronic pancreatitis[9]. Depletion of CTGF with neutralizing antibody significantly attenuated the pancreatic fibrosis in PL mice, suggesting that CTGF may be a potential therapeutic target for chronic pancreatitis.

The depletion of CTGF could not completely rescue the phenotype in PL mice, implying the existence of additional down-stream YAP1/TAZ mediators contributing to the fibrotic phenotype. Consistent with previous studies which showed that SSP1 is a potential YAP1/TAZ target, we found that it is upregulated in acinar cells soon after Lats1&2 deletion. SPP1 is a known mediator of inflammation and fibrogenesis in different organs[42],[43]. In the PL mice, SPP1 up-regulation co-localizes with YAP&TAZ-activated acinar cells and strongly correlates with activation of PSCs. Thus, SPP1 could be one of the YAP&TAZ effectors to contribute to the CP-like phenotype in PL mice. RNA-seq data showed that deletion of *Lat1&2* induced upregulation of several other known PSC activators, such as TGF-β, PDGF, CXCL12, and CXCL16, suggesting that the Hippo pathway may regulate the expression of multiple secreted factors to control acinar-PSC interactions.

Although our data support that PSCs could be directly activated by secreted factors from *Lats1&2* null acinar cells, many of these PSC activators are also immune regulators. Thus, our data do not exclude the possibility that the secreted factors released by *Lats1&2* null acinar cells may have direct effects on immune cells. Indeed, both activated PSCs and infiltrated immune cells are found in most early lesions in *Lats1&2* knockout mice, strongly supporting that the interactions between acinar cells and immune cells may contribute to the unique phenotype observed in our model. However, understanding the detailed mechanisms of this complex acinar-immune-PSC interaction is beyond the scope of this study and may require the development of new mouse models with appropriate immune backgrounds. In addition, we observed ADM after PSC activation and immune cell infiltration. Considering the well documented roles of inflammatory cytokines in the induction of ADM, it is unclear whether the ADM in Lats1&2 null pancreata is directly induced by activated YAP1/TAZ or by infiltrated immune cells. Additional mouse models are needed to answer this question.

In summary, a new mouse model has been established which exhibits pathological features similar to chronic pancreatitis. This model places the Hippo pathway impairment in acinar cells among the factors contributing to the pathogenesis of chronic pancreatitis. We reported the secretome alterations caused by *Lats1&2* deletion in adult acinar cells, highlighting the novel function of the Hippo pathway in regulating epithelial-stromal interaction in the adult pancreas. These data provide new insights into understanding the mechanisms of chronic pancreatitis progression and identify new potential therapeutic targets.

## Acknowledgements

The authors acknowledge Dr. Chris Wright and Dr. Matthias Hebrok for kindly providing the *Ptf1a^CreER^ Rosa26^LSL-YFP^* mouse line, Dr. Eric N. Olson for kindly providing the *Yap1^fl/fl^* and *Taz^fl/fl^* mouse line, and Kenneth Lipson of FibroGen, Inc. for providing CTFG neutralizing antibody. The authors thank Jake Leighton for helping with mouse breeding and genotyping. The authors thank Dr. Frank Gorelick (Yale University), Dr. Aida Habtezion (Stanford University), and Dr. Kenneth Lipson (FibroGen, Inc.) for their critical comments. The authors thank Dr. Minoti Apte (South Western Sydney Clinical School) and Dr. Mara Sherman (Oregon Health and Science University) and for sharing protocols and tips. The authors thank Dr. Zhao Lai and Mr. Huang-I Chen from UT Health San Antonio for assistance with RNA-sequencing and analysis. The authors thank Benjamin J. Daniel and Karla M. Gorena for their help with flow cytometry. Data was generated in the Flow Cytometry Shared Resource Facility, which is supported by UTHSCSA, NIH-NCI P30 CA054174-20 (CTRC at UTHSCSA) and UL1 TR001120 (CTSA grant).

## Transcript profiling

GEO accession code is GSE111640.

## Contributions

P.Wang, M. Gao and J. Liu designed and performed the experiments. F.Sharkey provided the histologic scoring of the experiments. P.Wang, M.Gao, J.Liu, and M.Nipper analyzed the data and wrote the manuscript. R.Johnson provided Lats1^fl/fl^ and Lats2^fl/fl^ mice. R.Johnson and H.Crawford provided reagents, tools, technical advice, and critical comments on manuscript.

