## supplemental data for "Deletion of Lats 1 and 2 in mouse pancreatic acinar cells induces pancreatic fibrosis and inflammation"

### Results

#### *Lats1&2* deletions do not induce autonomous proliferation, apoptosis, or ADM in adult acinar cells directly.

Pancreatic inflammation is frequently induced by acinar cell injury[1]. It is possible that loss of LATS1&2 resulted in autonomous cell injury by either promoting cellular proliferation or apoptosis. Cell injury may lead to the immune cell infiltration, PSC activation, and ADM which we observed in PL mice. The phenotypes in PL mice appeared very rapidly so it was hard to conclude whether or not the cell injury happened first. To investigate this possibility, we had to observe the cell behavior at the single-cell level. We subjected PL mice to a single injection of TAM at different doses (45 mg/kg, 90 mg/kg, and 180 mg/kg). PCR assays confirmed the occurrence of Cre-mediated *Lats1&2* deletions even four days after a single dose of 45 mg/kg TAM injection (Supplementary Figure 4A). We chose the 45 mg/kg dose for further experiments because it showed Cre-mediated recombination in individual acinar cells (Supplementary Figure 4B), as determined by YFP expression. Because one of the best known functions of LATS1&2 is to control cellular proliferation, we first tested whether *Lats1&2* deletion induced the proliferation of acinar cells. We examined Ki67 expression in the pancreata of PL mice at 4, 8, and 12 days after a single TAM injection (45 mg/kg) (Figure 2A). No $YFP^+$ cells expressed Ki67 at every time point. We then co-stained YFP with cleaved-caspase 3 in these pancreata to test whether these cells underwent apoptosis. The results showed no YFP+ cells that were co-stained with cleaved-caspase3 (Figure 2B). In addition, we quantified the number of $YFP^+$ cells in these mice and found no significant differences during the 12-day period (Figure 2C). IHC assay revealed YAP1 and TAZ nuclear translocation in PL pancreas, further supporting the successful deletion of *Lat1&2* (Figure 2D). Collectively, these data indicated that loss of *Lats1&2* did not affect adult acinar cellular proliferation or apoptosis directly.

Recent studies suggested that inactivation of the Hippo pathway plays important roles in promoting ADM. Thus, we tested whether loss of *Lats1&2* could induce ADM in adult acinar cells. We did not observe YFP positive cells expressing CK19, indicating that no ADM occurred in *Lats1&2* null acinar cells (Figure 2B). Taken together, these data suggest that the pancreatitis-like phenotype of PL mouse was unlikely to be initiated by proliferation, apoptosis, or ADM.

### Supplementary materials and methods

#### *Generation of conditional knockout mice*

We generated (1) *Ptf1a*$^{CRE\text{-}ER}$*Rosa26*$^{LSL\text{-}YFP}$ mice (P mice, control), (2) *Ptf1a*$^{CRE\text{-}ER}$*Rosa26*$^{LSL\text{-}YFP}$*Lats1*$^{fl/fl}$*Lats2*$^{fl/fl}$ *mice* (PL mice), (3) *Ptf1a*$^{CRE\text{-}ER}$*Yap1*$^{fl/fl}$*Taz*$^{fl/fl}$ mice (PTY mice), and (4) *Ptf1a*$^{CRE\text{-}ER}$*Rosa26*$^{LSL\text{-}YFP}$*Lats1*$^{fl/f}$*lLats2*$^{fl/fl}$*Yap1*$^{fl/fl}$*Taz*$^{fl/fl}$ mice (PLTY mice). All offspring were genotyped by PCR of genomic DNA from the toe with primers specific for the *Ptf1a*$^{Cre\text{-}ER}$, *Rosa26*$^{LSL\text{-}YFP}$, *Lats1*, *Lats2*, *Yap1*, and *Taz* transgenes. To induce the conditional knockout, 6- to 12- week old mice were treated with 180mg/kg of tamoxifen (TAM, Sigma-Aldrich, T5648-5G) which was dissolved in corn oil (Sigma-Aldrich, C8267) via intraperitoneal (i.p) injection once every 24 hours for five consecutive days. PCR was used for validation of knockout alleles (Supplementary Table 2).

#### *Caerulein-induced acute and chronic pancreatitis*

Caerulein (American Peptide Company, 46-1-50) was used to induce both acute and chronic pancreatitis mouse models. For acute pancreatitis (AP), caerulein was administered through eight intraperitoneal (i.p.) injections (50μg/kg) at hourly intervals per day for two days[2]. Pancreatic inflammation was assessed by H&E staining on Days 1, 2, 3, 4 and 6 after induction of acute pancreatitis. For chronic pancreatitis, mice were subjected to repetitive episodes of AP

(50ug/kg, eight injections at hourly intervals), every other day for four weeks. Mice were sacrificed four weeks after the initial caerulein injection.

***Tissue preparation***

Mice were euthanized and pancreata were dissected into ice-cold phosphate-buffered saline (PBS). For histological examination, mice pancreata were fixed in 4% Paraformaldehyde (PFA) overnight at $4^oC$ and then processed for paraffin or O.C.T embedding. Sections were cut at 5 μm and then used for hematoxylin and eosin (H&E) staining or immunostaining.

***Immunostaining***

Primary and secondary antibodies used in this study are listed in supplementary Table S2. For immunohistochemistry staining (IHC), tissues were deparaffinized, rehydrated, and immersed and heated in a citrate-based antigen unmasking solution (Vector, H300) for 30 minutes. Endogenous peroxidase activity was quenched with 3% hydrogen peroxide in $ddH_2O$. Sections were blocked with 5% donkey serum in 0.1%TBST (Tween®20, Sigma-Aldrich, P1379-500ml) for one hour at room temperature, and then incubated with primary antibody at $4^oC$ overnight. Biotin-labeled secondary antibodies (1:250, BD Pharmingen™) and Streptavidin-Horseradish peroxidase (SAv-HRP, BD Pharmingen™ 550946) were applied to amplify the target antigen signal, then sections were developed with DAB substrate (Dako, K3468), followed by counterstaining with Hematoxylin (Biocare Medical). Finally, sections were covered with poly-Mount medium (Polysciences, Inc. Cat # 08381). For immunofluorescence staining, frozen sections were blocked with 10% donkey serum plus 1% bovine serum albumin (BSA) in PBST (0.025% Triton X-100, Fisher Scientific. CAS9002-93-1) for one hour at room temperature. Sections were incubated with primary antibodies at $4^oC$ overnight followed by incubation with the Alexa Fluor® secondary antibodies (1:250, Jackson ImmunoResearch) for one hour at room temperature. Sections were covered with a drop of ProLong Gold anti-fade reagent with DAPI

(Invitrogen, P36935) for observation. All Images were obtained with a Microsystems DMI6000 B microscope and microscope software (Leica Microsystems).

***Western blot analysis***

After the treatments described in the text and figure legends, proteins were isolated from the tail of the pancreas. Briefly, pancreata were homogenized in RIPA lysis buffer containing phosphatase and protease Inhibitors (GenDEPOT, P3100-001 and P3200-001) on ice. After 10 cycle of sonication (Bioruptor® Pico Sonication System, Cat.No.B01060001), proteins were extracted and the concentration was determined by a BCA kit (Pierce, 23228). Equal amounts of 50ug protein were resolved in SDS-PAGE, then transferred to PVDF membrane (ThermoFisher Scientific, Prod # 88520) and blocked with 5% milk in PBST (0.1% Tween® 20, Sigma-Aldrich, P1379-500ml) for one hour at room temperature. Membranes were incubated with primary antibodies at 4$^{o}$C overnight and then exposed to secondary antibodies for one hour. Finally, protein expression was determined by treating the membrane with Clarity™ Western ECL substrate (Bio-Rad, Cat. 1705060) and measuring luminescence with an Amersham Imager 600 system (GE Healthcare). (All antibody information and working concentrations are shown in Supplementary Table 1).

***Reverse transcription (RT) and real-time PCR analysis***

Total RNA was extracted from the tail of the pancreas using a lysis binding buffer (Ambion, Cat. AM1560) and Direct-zol RNA miniprep kit (Zymo Research, R2052). The RNA concentration was measured using Nanodrop2000 (ThermoFisher Scientific). 400ng of RNA was subjected to reverse transcription by using a high capacity cDNA reverse transcription kit (Applied Biosystems, 4368814), following the manufacturer's protocol. The cDNA samples were amplified with iTaq Universal SYBR Green Supermix (Bio-Rad, 172-5271) in a CFX96 Real-Time system (Bio-Rad). GAPDH was used as endogenous control for normalization and the relative expression of mRNA was calculated based on the $^{\Delta\Delta}$Ct method. The information of the primers is available in Supplementary Table 3.

***Flow cytometry analysis***

Single-cell solutions from P and PL mice were prepared according to previous reports. Briefly, six-week aged mice P and PL were treated with 180mg/kg of tamoxifen for five consecutive days. Four days later, mice were anesthetized, and then perfused with cold PBS. Pancreas was digested by collagenase (Sigma-Aldrich, C9891-1G), then cells eventually filtrated through 5ml polystyrene round-bottom tube with cell-strainer cap (Corning, #352235). All antibodies used for flow cytometry were purchased from Biolegend: APC-CD45 (103111, 1:100), Percp/Cy5.5-CD11b (101228, 1:100), BV421-F4/80 (123137, 1:100). Cells were incubated with the antibodies in FACS buffer at 4$^{o}$C for 15 mins. After incubation, cells were washed once and resuspended into FACS buffer. Sorting experiments were conducted on BD FACSAria with BD FAC SDiva software.

***Isolation and culture of mouse PSCs***

Primary mouse PSCs were isolated using a previously reported gradient centrifugation method with a few modifications[3]. Briefly, pancreata were dissected from wild type mice at the ages of 4-6 weeks. After all adipose tissues were carefully trimmed off, pancreata were cut into small pieces in cold HBSS buffer. Tissues from 2 mice were transferred into a 50ml tube containing 10ml HBSS buffer (0.05% pronase, 0.035% collagenase P, 0.1% DNAse, 10mM HEPES, and 0.01 Trypsin inhibitor). Tissues were incubated in a 37 °C water bath for 10 minutes and then dissociated by pipetting up and down with a 10 ml tip 10 times. The tissues were incubated in a 37 °C water bath for an additional 5-10 minutes and dispersed by pipetting gently up and down with a 10 ml pipette 10 times. The cell suspension was filtered through a 70 um cell strainer. Cells were washed twice and suspended in 2ml HBSS buffer containing 0.3%

BSA. 4ml 60% Optiprep was added to the cell suspension and mixed well in a 15ml tube. 2ml HBSS buffer was carefully added to the top of the Optiprep gradient. PSCs were separated by centrifugation at 1400Xg for 20min. PSCs were suspended in DMEM:F-12 medium containing 10%FBS, 15mM HEPES, 4Mm glutamine, and antibiotics (penicillin 100U/ml and streptomycin 100 ug/ml). Cells were seeded on a 10 well microscope slide (6000 cells/well) and treated with CTGF and SPP1 for 3 days.

***RNA-sequencing and data analysis***

Total RNA was isolated using RNA lysis binding buffer (Ambion, Cat. AM1560) at Days 1, 2, and 3, as well as control condition. RNA quality was assessed by Bioanalyzer and poly A(+) RNA was isolated by oligo-dT purification and fragmented using divalent cations under elevated temperature. cDNA fragment libraries were synthesized following the TruSeq mRNA-seq Library Preparation protocol (Illumina). For total of 16 samples, we obtained about 31.4 million sequence reads per sample, using Illumina HiSeq system at the Greehey Children's Cancer Research Institute Genome Sequencing Facility (GCCRI GSF), utilizing a 50bp single-read sequencing protocol.

Short sequence reads were then aligned with TopHat[4] to mouse genome (UCSC mm9), allowing no more than two mismatches in the alignment. Expression abundance of each gene was evaluated with read count and RPKM (read per kilobase of transcript per million reads mapped). Differential gene expression was calculated using DESeq[5] to obtain fold-change, p-value, and adjusted p-value by Benjamini-Hochberg correction for multiple tests. Differentially expressed genes were selected based on the following criteria: 1) fold-change > 2, 2) adjusted p-value < 0.05, and 3) RPKM > 1.  Using the significance criterion and for comparison between Days 1, 2, and 3 to control, we obtained 40, 405 and 765 differentially expressed genes (DEGs), respectively. To access the overall expression change across time, a heatmap was created using all DEGs from three comparisons. We also performed K-means algorithm (MATLAB,

Mathworks) with all DEGs into 9 clusters and then merged upregulated and downregulated genes into two representative clusters (300 and 232 genes, respectively). Biological functional assessment of these differentially expressed genes was performed by using Database for Annotation, Visualization and Integrated Discovery (DAVID, http://david.abcc.ncifcrf.gov/), with top 10 most enriched Biological Processes (BP) were provided in the figure.

### Supplementary Tables

1. List of antibodies used in this study

| Primary Antibody | Catalog | Dilution | Company | Application |
|---|---|---|---|---|
| Lats1 (C66B5) | 3477 | 1:1000 | Cell signaling technology | WB |
| Last2 | A300-479A | 1:1000 | Bethyl Laboratories,INC. | WB |
| YAP1 | 13584-1-AP | 1:1000 | Proteintech$^{TM}$ | WB |

| | | | | |
|---|---|---|---|---|
| P-YAP1(Ser127) | 4911S | 1:1000 | Cell signaling technology | WB |
| TAZ | 23306-1-AP | 1:1000 | Proteintech™ | WB |
| CTGF (I-20) | sc-14939 | 1:1000 | Santa Cruz Biotechnology | WB |
| γ-H2AX | 2577S | 1:1000 | Cell signaling technology | WB |
| GAPDH | sc-32233 | 1:200 | Santa Cruz Biotechnology | WB |
| Tubulin | 11224-1-AP | 1:1000 | Proteintech™ | WB |
| YAP1(D8H1X) | 14074 | 1:200 | Cell signaling technology | IHC |
| TAZ | HPA007415-100UL | 1:600 | Sigma-Aldrich | IHC |
| F4/80 [CI:A3-1 clone] | Ab6640 | 1:100 | Abcam | IHC |
| SPP1 | AF808 | 1:200 | R&D | IHC, IF |
| CK19 | TROMA-III | 1:50 | DSHB | IHC, IF |
| CD45 (Clone 30-F11) | 70-0451 | 1:100 | Tonbo Biosciences | IHC, IF |
| α-SMA | 14-9760-80 | 1:3000 | eBioscience | IHC, IF |
| Anti-GFP | GFP-1020 | 1:500 | AVES LABS,INC. | IF |
| Amylase (C-20) | sc-12821 | 1:100 | Santa Cruz Biotechnology | IF |
| Collagen I | ab21286 | 1:100 | Abcam | IF |
| Ki67 | RM-9106-S | 1:100 | Thermo Scientific | IF |
| Activated Caspase 3 | G748A | 1:250 | Progema | IF |
| APC-CD45 | 103111 | 1:100 | Biolegend | FC |
| BV421-F4/80 | 123137 | 1:100 | Biolegend | FC |
| Percp/Cy5.5-CD11b | 101228 | 1:100 | Biolegend | FC |

2. Primer sequences used for genotyping

| Lats1 (F1+R: WT-350bp; F2+R: KO-230bp) | Lats2 (F1+R: WT-300bp; F2+R: KO-250bp) |
|---|---|
| F1: TTGTTGCTGGTGTTGTTT CC | F1: GCGCATGCCTTTAATCCTAGC |
| F2: AGGATGTAGTGAAGGCGTGTAAC | F2: CTATCGCTAGGCTGTTCCCAC |
| R: AGACCTCGTCGCACAGAATG | R: CTGAGCAACGACTCCAGGAAC |
| YAP1 (F1+R1: WT-457bp; F1+R2: KO-338bp) | TAZ (F1+R1: WT-496bp; F1+R2: KO-704bp) |
| F1:ACATGTAGGTCTGCATGCCAGAGGAGG | F1: GGCTTGTGACAAAGAACCTGGGGCTATCTGAG |
| R1:AGGCTGAGACAGGAGGATCTCTGTGAG | R1: CCCACAGTTAAATGCTTCTCCCAAGACTGGG |
| R2:TGGTTGAGACAGCGTGCACTATGGAGC | R2: AACTGCTAACGTCTCCTGCCCCTGACCTCTC |

3. Primer sequences used for Real-time PCR in this study

| Gene name | Forward | Reverse |
|---|---|---|
| Lats1 | GCGATGTCTAGCCCATTCTC | GGTTGTCCCACCAACATTTC |
| Lats2 | AGCCTGACAACATACTCATCG | AATCCAGTGCAGAGGCCAAA |
| YAP1 | TACTGATGCAGGTACTGCGG | TCAGGGATCTCAAAGGAGGAC |
| TAZ | GAAGGTGATGAATCAGCCTCTG | GTTCTGAGTCGGGTGGTTCTG |
| CTGF | GGCCTCTTCTGCGATTTCG | GCAGCTTGACCCTTCTCGG |
| SPP1 | AGCAAGAAACTCTTCCAAGCAA | GTGAGATTCGTCAGATTCATCCG |
| CXCL12 | TGCATCAGTGACGGTAAACCA | TTCTTCAGCCGTGCAACAATC |
| CXCL16 | CCTTGTCTCTTGCGTTCTTCC | TCCAAAGTACCCTGCGGTATC |
| TGF-β1 | CCACCTGCAAGACCATCGAC | CTGGCGAGCCTTAGTTTGGAC |
| TGF-β2 | CTTCGACGTGACAGACGCT | GCAGGGGCAGTGTAAACTTATT |
| TGF-β3 | GGACTTCGGCCACATCAAGAA | TAGGGGACGTGGGTCATCAC |

| | | |
|---|---|---|
| PDGF | GAGGAAGCCGAGATACCCC | TGCTGTGGATCTGACTTCGAG |
| YM1 | TGGTGAAGGAAATGCGTAAA | GTCAATGATTCCTGCTCCTG |
| IL-10 | GCTGGACAACATACTGCTAACC | ATTTCCGATAAGGCTTGGCAA |
| CCL17 | TACCATGAGGTCACTTCAGATGC | GCACTCTCGGCCTACATTGG |
| TNF-$\alpha$ | CCAAAGGGATGAGAAGTTCC | CTCCACTTGGTGGTTTGCTA |
| SOCS1 | CTGCGGCTTCTATTGGGGAC | AAAAGGCAGTCGAAGGTCTCG |
| NOS2 | GTTCTCAGCCCAACAATACAAGA | GTGGACGGGTCGATGTCAC |
| GAPDH | AGGTCGGTGTGAACGGATTTG | GGGGTCGTTGATGGCAACA |

### Supplementary Figure Legends

Supplementary Figure 1: Hippo pathway is dynamically regulated in pancreatitis models. (A) Quantification of Western blot of LATS1, LATS2, P-YAP1, YAP1, and TAZ in untreated and AP mice. Untreated mice served as the control group. Tubulin was used as the internal control (n=4); (B) Immunochemistry Staining to detect YAP1 expression on day 1 after AP induction (n=4); (C) Quantification of Western blot of LATS1, LATS2, YAP1, and TAZ in untreated and CP mice. Untreated mice served as the control group. Tubulin was used as the internal control (n=4). **, P<0.01.

Supplementary Figure 2: Acinar-specific Hippo pathway inactivation induced pancreatic inflammation-associated phenotypes in mice. (A) Mice breeding strategy and experimental design; (B) Time course analysis for body weight changes of P and PL mice after 5 consecutive TAM injections (n=5); (C) Body weight and blood glucose levels in P and PL mice on day 10 after final TAM injection; (D) Time course H&E analysis for pancreata of PL mice with 5 consecutive TAM injections.

Supplementary Figure 3: YAP and TAZ activations were regulated by Lats1&2 in adult acinar cells. (A) Scheme of the mouse *Lats1* and *Lats2* locus and the strategy of detecting *Lats1&2* deletions. Confirmation of the excisions of *Lats1* exon 4 (Deletion: 230bp) and *Lats2* exon 5 (Deletion: 250bp) in PL mouse by PCR. The primer sequences are indicated as P1, P2, P3 for *Lats1* detection, and P1’, P2’, P3’ for *Lats2* detection; (B) *Lats1&2* deletion induced YAP1&TAZ translocation was detected by IHC staining in P, PL, PL1KO, and PL2KO mice. n=6.)

Supplementary Figure 4: Acinar-specific Lats1&2 depletions induced pancreatitis-associated histological alterations. (A) ADM was quantified by counting YFP and CK19 double positive cell numbers. CD45 and $\alpha$SMA were quantified by IHC profiler score (n=5). *, P<0.05, **, P<0.01. Representative immunofluorescence staining with (B) anti-YFP (Green), anti-CK19 (Red), anti-Ki67 (White) antibodies and with (C) anti-YFP (Green), anti-CK19 (Red), anti-Cleaved-Caspase3(White) antibodies in P and PL pancreata. Nuclei stained with DAPI (Blue). Ki67 and Cleaved-Caspase3 were quantified by IHC profiler score (n=5); **, P<0.01.

.

Supplementary Figure 5. Mosaic Lats1&2 deletion induced long-lasting pancreatic inflammation. (A) PL mice were injected once with 45mg/kg, 90mg/kg, or 180mg/kg of TAM, respectively. Confirmation of the excisions of *Lats1* exon 4 and *Lats2* exon 5 by PCR at 45mg/kg of TAM condition. *Lats1* deletion: 230bp; *Lats2* deletion: 250bp. (B) Anti-YFP antibody (Green) was used to stain *Lats1&2* null cells 2 days later. Nuclei stained with DAPI (Blue). (C) 3 weeks later, mice among injection groups were euthanized and pancreata were stained with H&E, anti-CD45, anti-$\alpha$SMA, and anti-CK19 antibodies (n=4).

Supplementary Figure 6: Loss of Lats1&2 does not affect acinar cell proliferation, apoptosis, or ADM directly. (A) Schematic diagram of experimental design (n=3). (B) PL mice were injected once with 45mg/kg of TAM. Cell proliferation, apoptosis, and ADM were analyzed on D4, D8, and D12 by co-staining anti-YFP with anti-Ki67, anti-cleaved-caspase 3, anti-amylase, and anti-CK19 antibodies. Nuclei stained with DAPI (Blue). Percentages of Ki67, Cleaved-Caspase3 and YFP cells were quantified by IHC profiler score. (D) YAP1 and TAZ were detected by IHC in PL mice after injection with 45mg/kg of TAM.

Supplementary Figure 7: Effect of Lats1&2 knockout on ADM, PSC activation, and immune cell infiltration. (A)Time course quantification of ADM, PSC activation, and immune cell infiltration in the pancreas of PL mice after a single dose TAM injection (180mg/kg) (n=4). (B) PL mice were injected once with 180mg/kg of TAM. ADM, PSC activation, and immune cell infiltration were detected by anti-CK19, anti-αSMA, and anti-CD45 antibodies on Day 10 and Day 20 after TAM injection. (C) P and PL mice were consecutively injected with 5 doses (180mg/kg) of TAM. Primary pancreatic acini were isolated 3 days after final injection and embedded into collagen for 3D culture (n=3). Cells were treated with or without TGFα (100ng/mL) for 5 days.

Supplementary Figure 8. Generation of mice with *Lats1*, *Lats2*, *Yap1*, *Taz* quadruple deletions in pancreatic acinar cells. (A) Generation of PTY mice and the strategy for detecting *Yap1*&*Taz* deletion. H&E staining were performed in P and PTY mice; (B) PLTY mice breeding strategy and experimental design; (C) Quantification of Western blot of LATS1, LATS2, YAP1, and TAZ in PL and PLTY mice. P mice served as the control group. Tubulin was used as the internal control (n=6); **, P<0.01. (D) Quantification of αSMA and CD45 immunostaining in P, PL and PLTY mice showed 90.1% reduction of inflammation and 95.2% reduction of stromal reaction in PLTY mice compared with PL mice. ADM was quantified by counting YFP and CK19 double positive cell numbers. There was an 84.5% reduction of ADM in PLTY mice compared to PL mice (n=6). **, P<0.01.

Supplementary Figure 9: Depletion of CTGF with neutralization antibody to rescue Lats1&2 knockout-induced phenotypes. (A) Quantification of Western blot of CTGF in PL and PLTY mice. Tubulin was used as the internal control (n=6); **, P<0.01. (B) Quantitative analysis of FG-3019 treatment effects on Lats1&2 deletion-induced ADM, PSC activation, and immune cell infiltration in mouse pancreas (n=6); *, P<0.05. (C) Effects of FG-3019 treatment on body weights of PL mice that received TAM injection. Mice injected with FG-huIgG antibody were used as the control (n=5-6).

Supplementary Figure 10 Lats1&2 deletions in pancreatic acinar cells induce chronic pancreatitis-like phenotype rapidly and SPP1 is strongly associated with PSC activation. (A) H&E staining of PL mice after TAM injection of 180mg/kg/day for 5 consecutive days via i.p. n=4. (B) αSMA, CK19 and CD45 IHC staining in consecutive sections at D2 and D3 after final injection. (C) The mRNA expression of Lats1, Lats2, Ctgf, Cyr61, and Spp1 were measured by Q-PCR in P and PL (D2) mice. **, $p<0.01$. (D) Co-stain of αSMA (Red) with SPP1 (Green) in P and PL (D2 and D3) mice by Immunofluorescence. (E) Small lesion was co-stained with αSMA (Red) and SPP1 (Green) in PL mice (180mg/kg of TAM, D10) by Immunofluorescence. Nuclei stained with DAPI (Blue).

Supplementary Figure 11. Examination of the effects of CTGF and SPP1 on PSC activation *in vitro*. (A) Representative immunofluorescent staining of GFAP in isolated mouse PSCs. (B) Quantitative analysis for αSMA and Edu incorporation in CTGF or SPP1-stimulated cells (n=3). *, $P<0.05$.

Supplementary Figure 12. Examine the effects of Lats1&2 deletions on macrophage polarizations. (A) Time course analysis of immune cell infiltration in the pancreas of P and PL mice after 5 consecutive TAM injections. Immune cells were stained with anti-CD45 antibody (n=3). (B) Gating strategy to sort macrophages for quantitative RT-PCR assay. Immune cells were stained with CD45 (P1: red). CD45+CD11b+F4/80+ macrophages were sorted (P2: blue)

Supplementary Figure 1

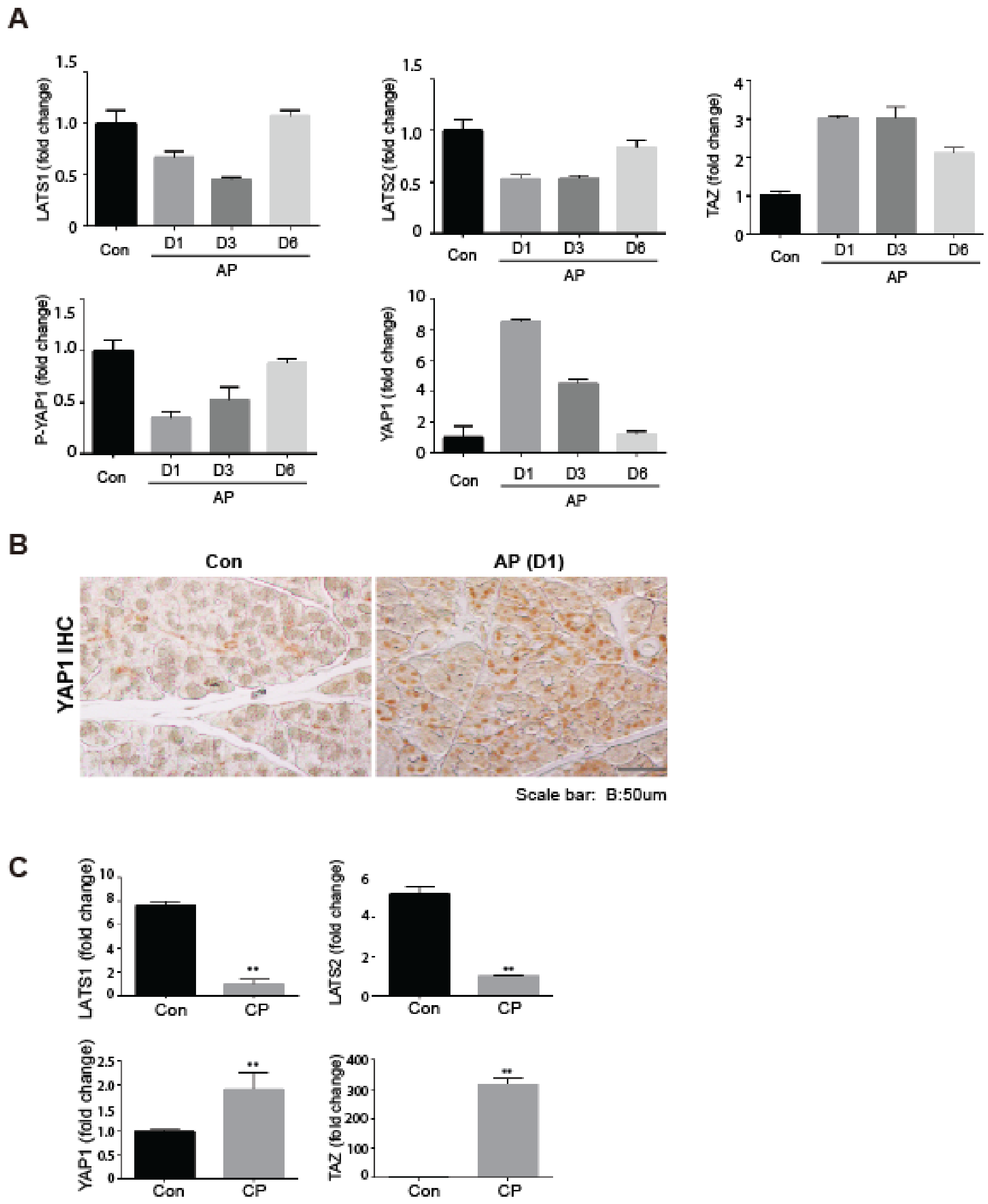

Supplementary Figure 2

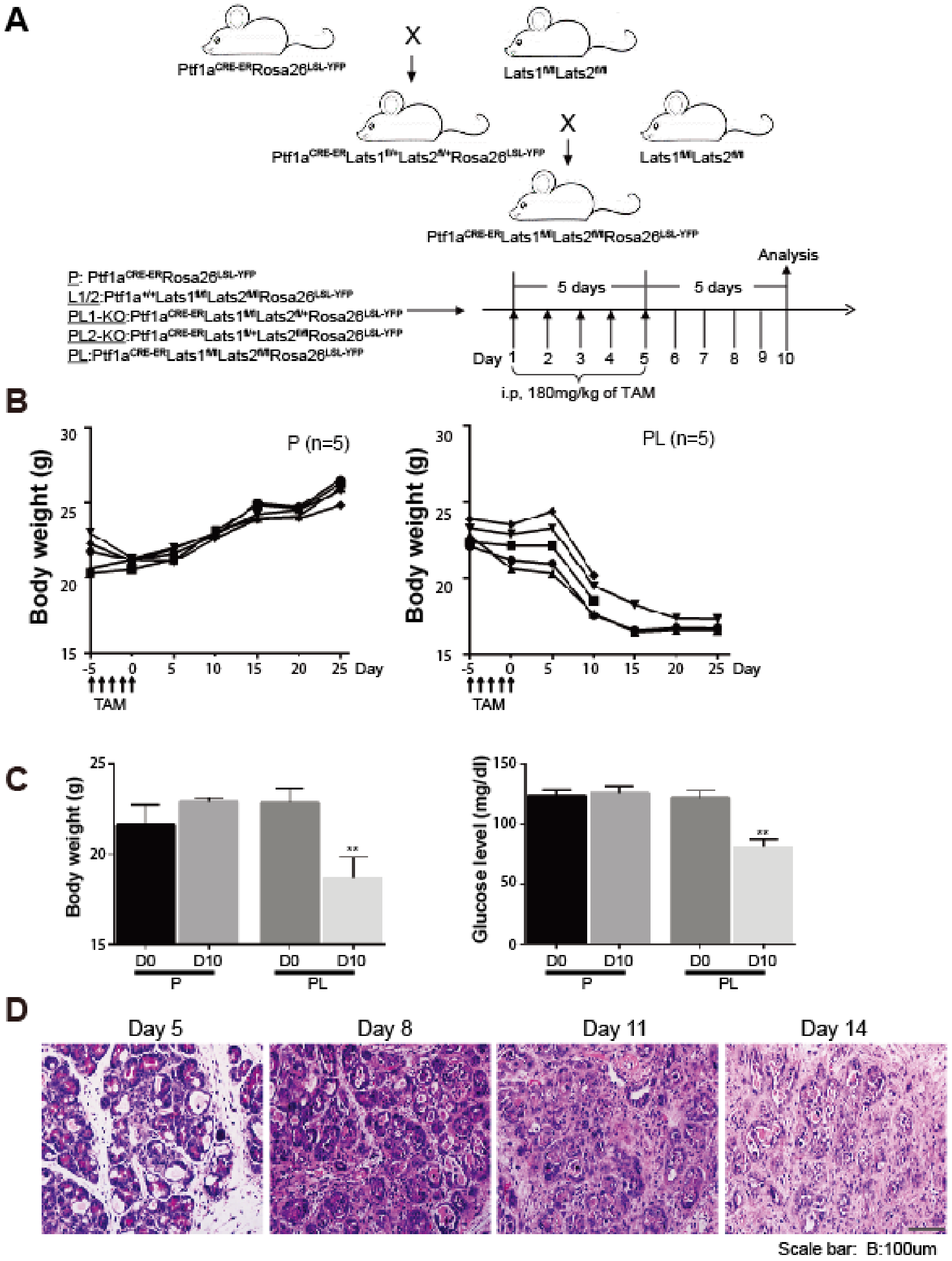

Supplementary Figure 3

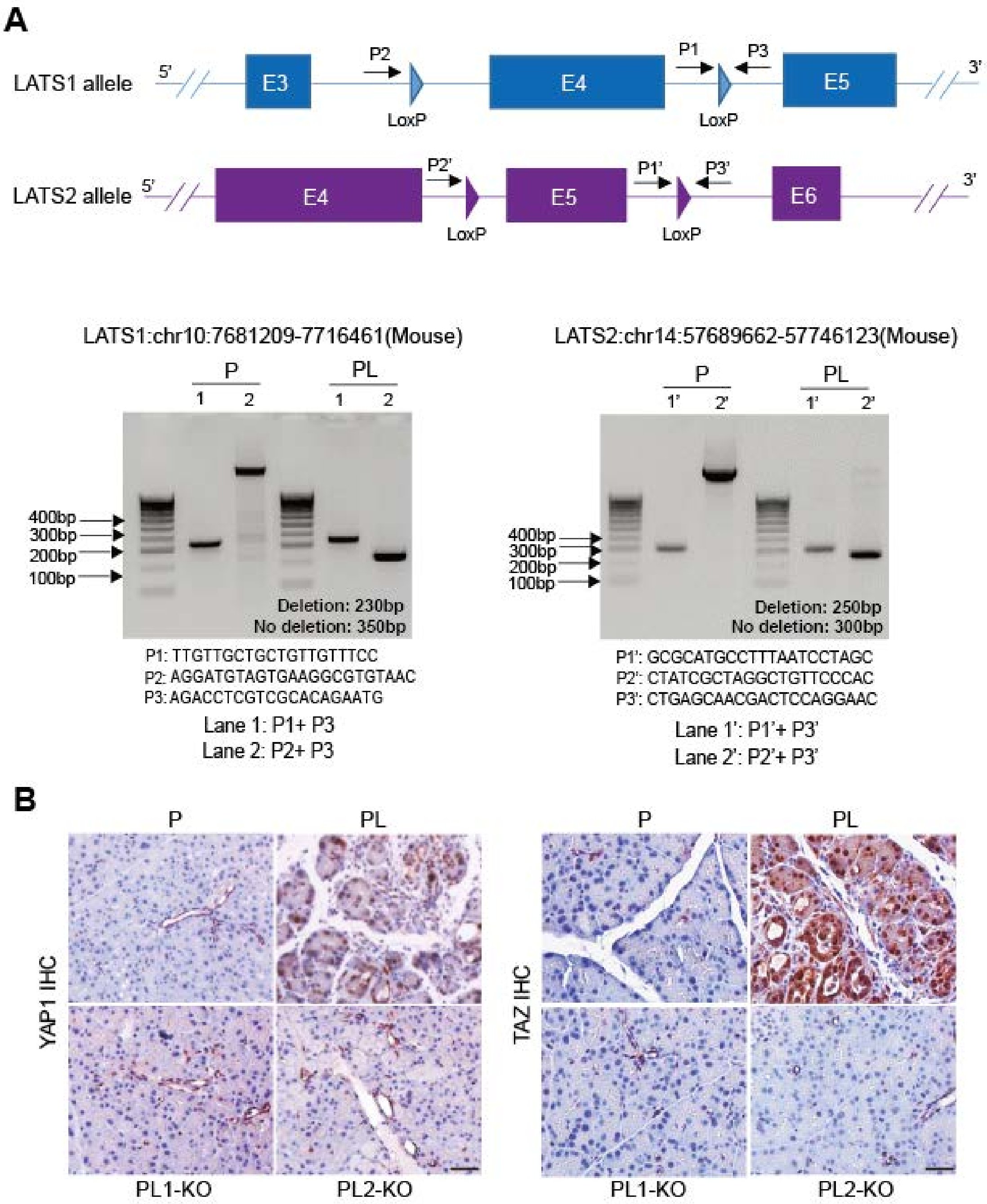

Supplementary Figure 4

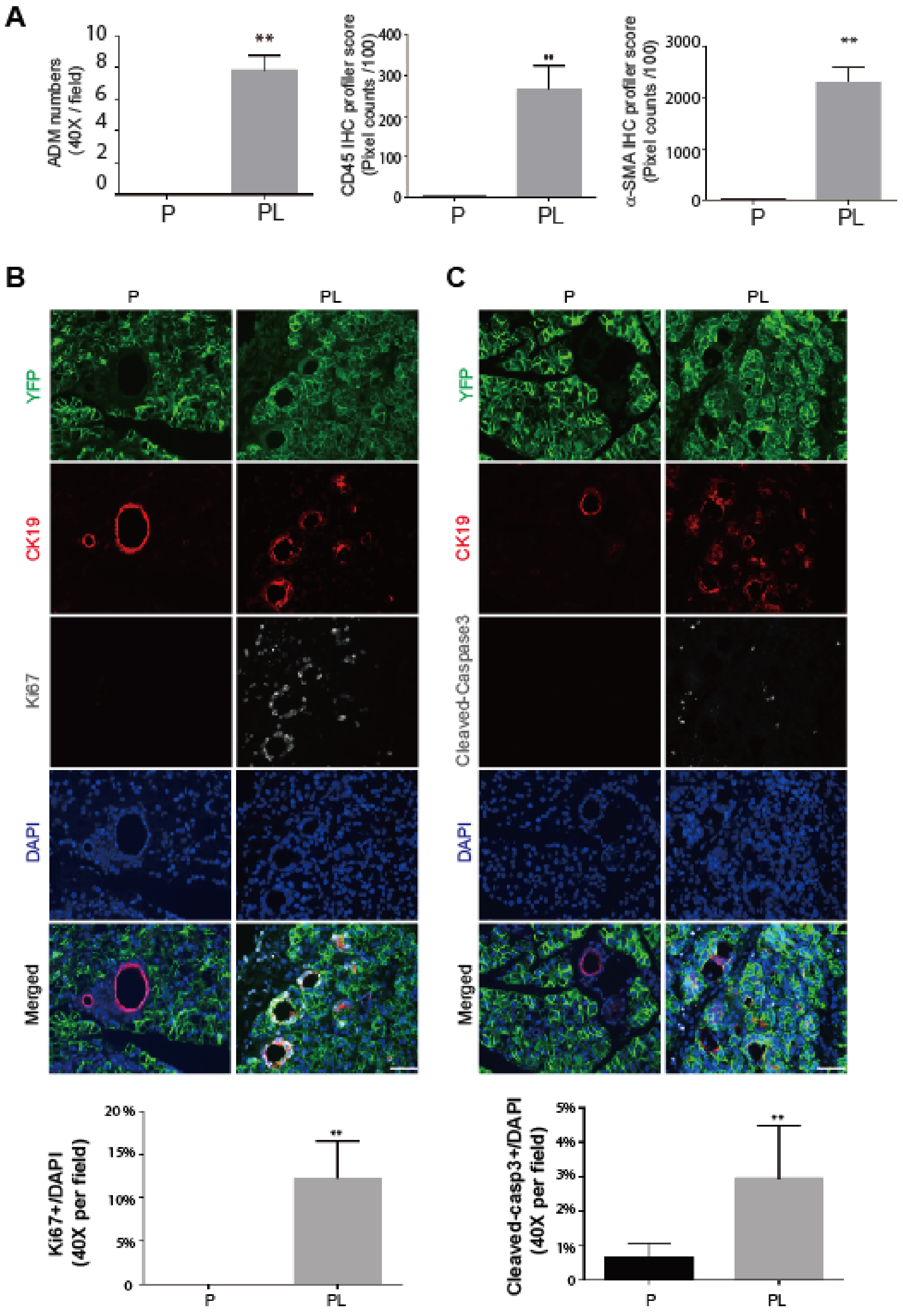

Supplementary Figure 5

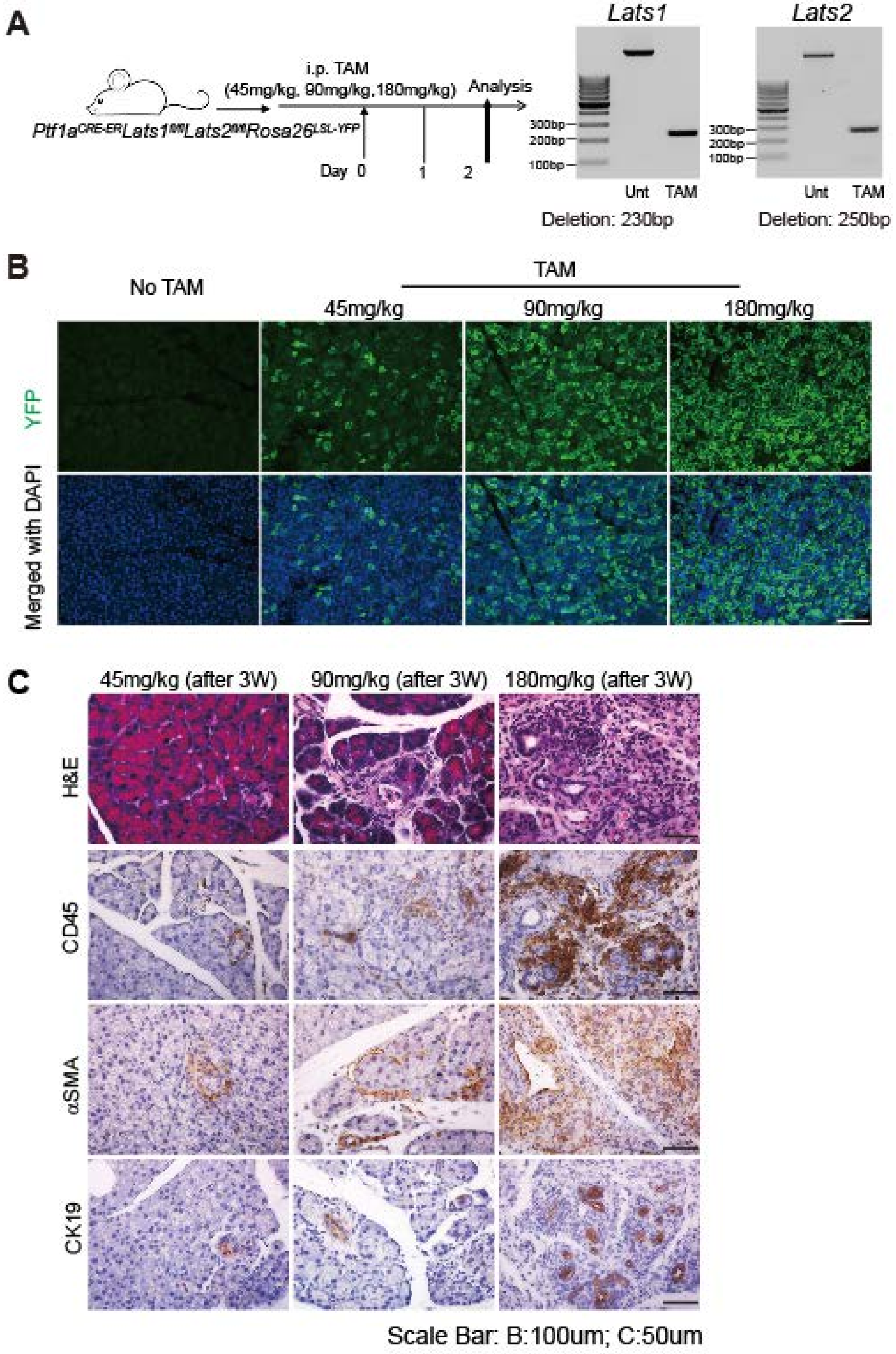

Supplementary Figure 6

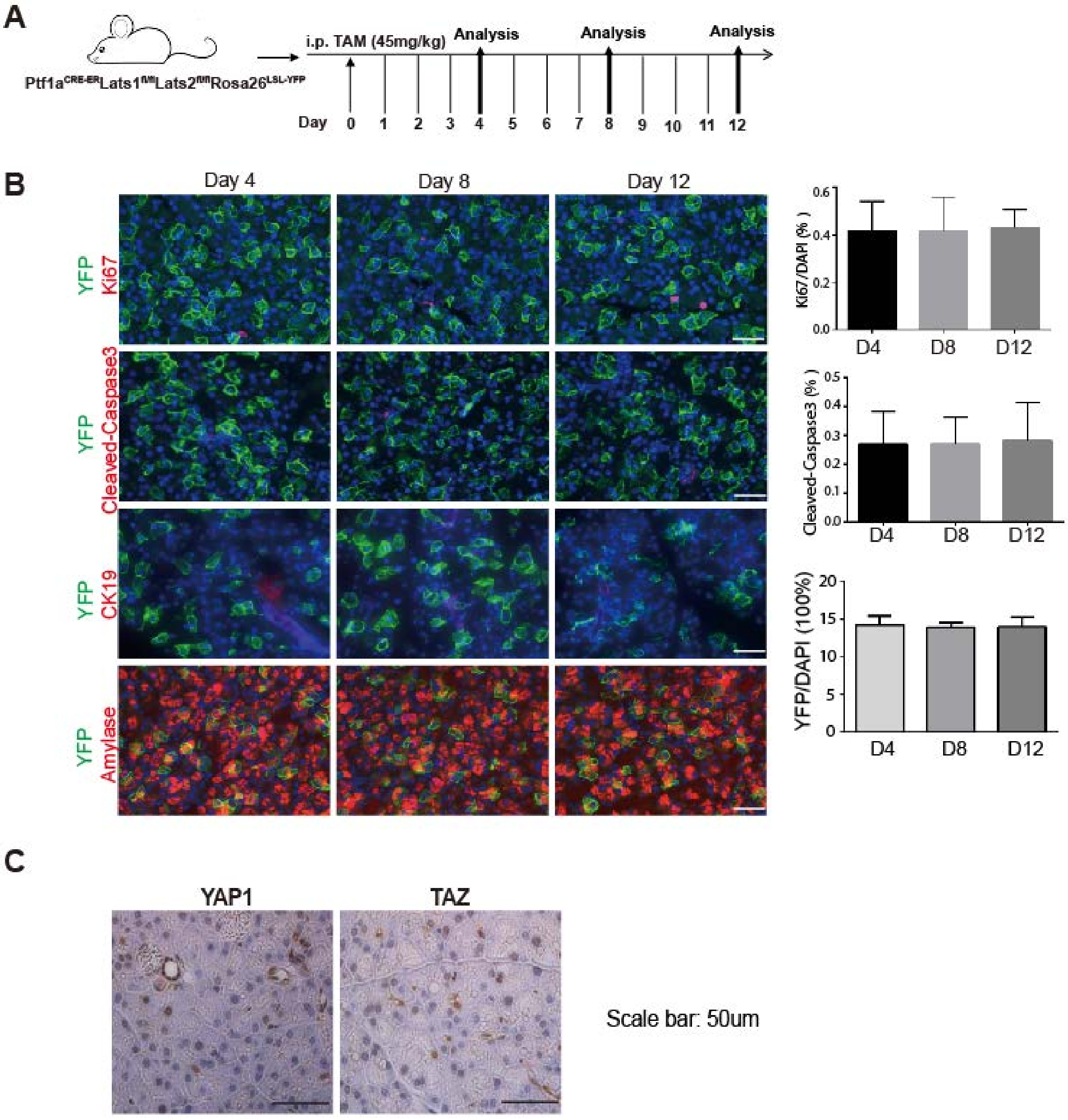

Supplementary Figure 7

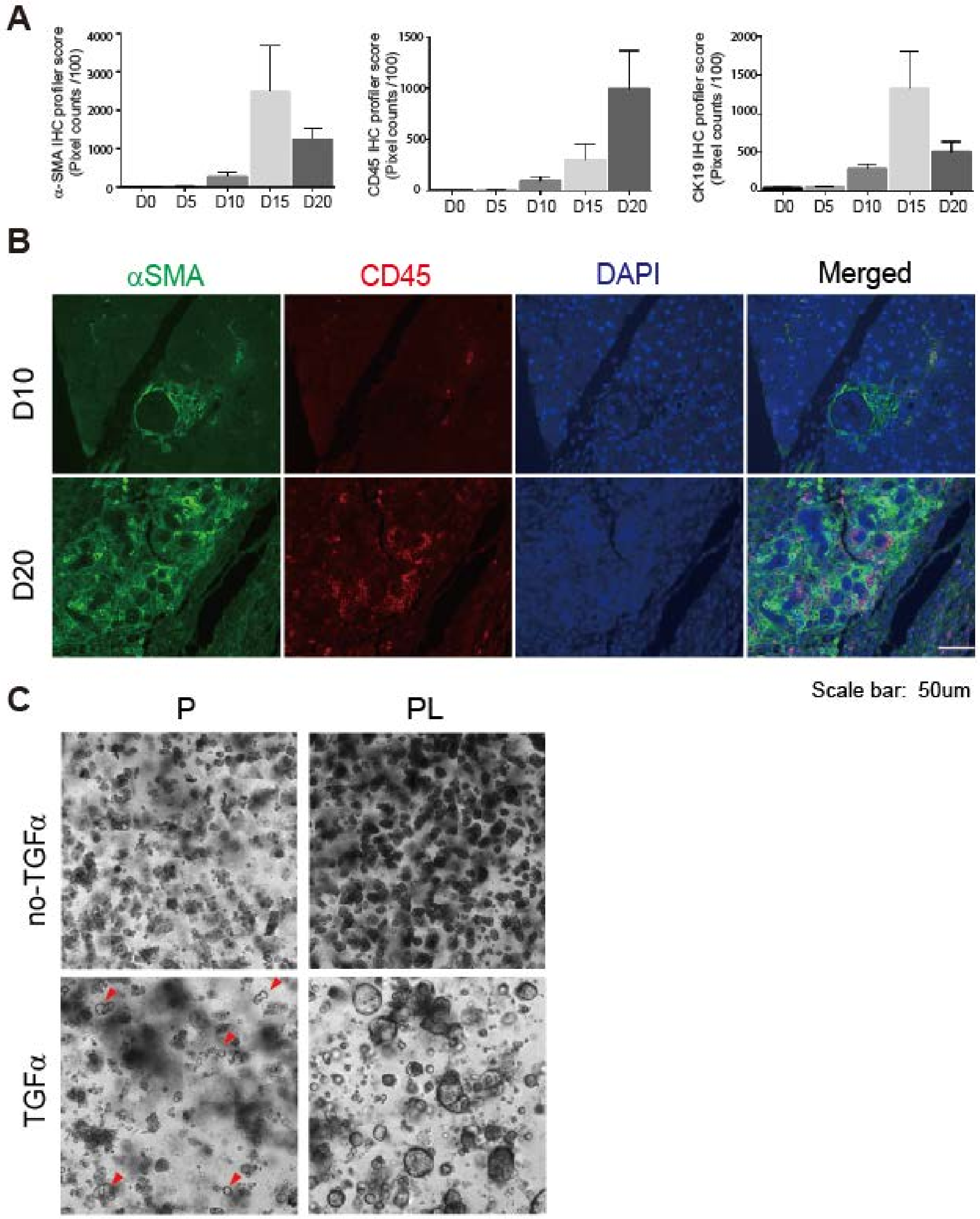

Supplementary Figure 8

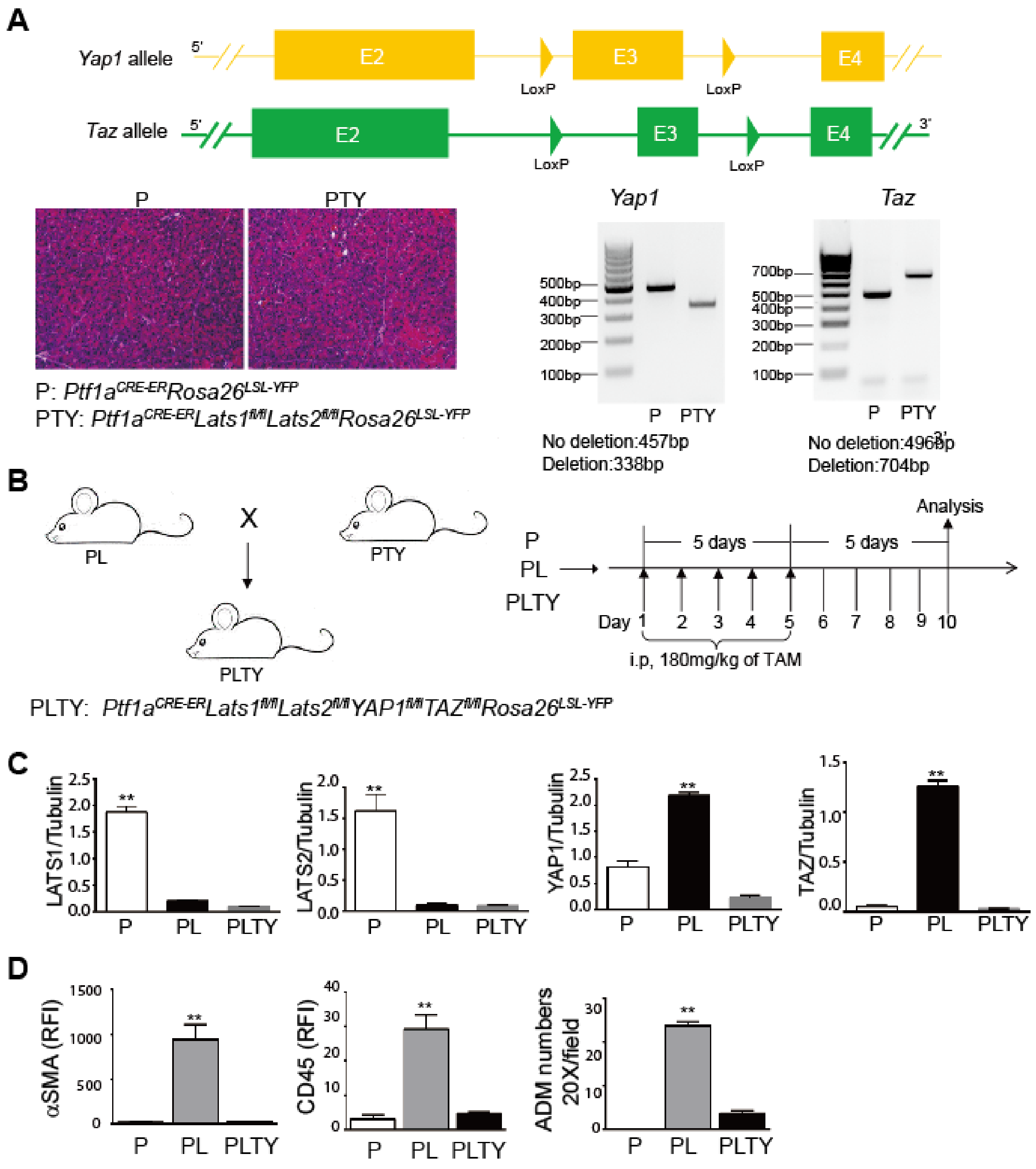

Supplementary Figure 9

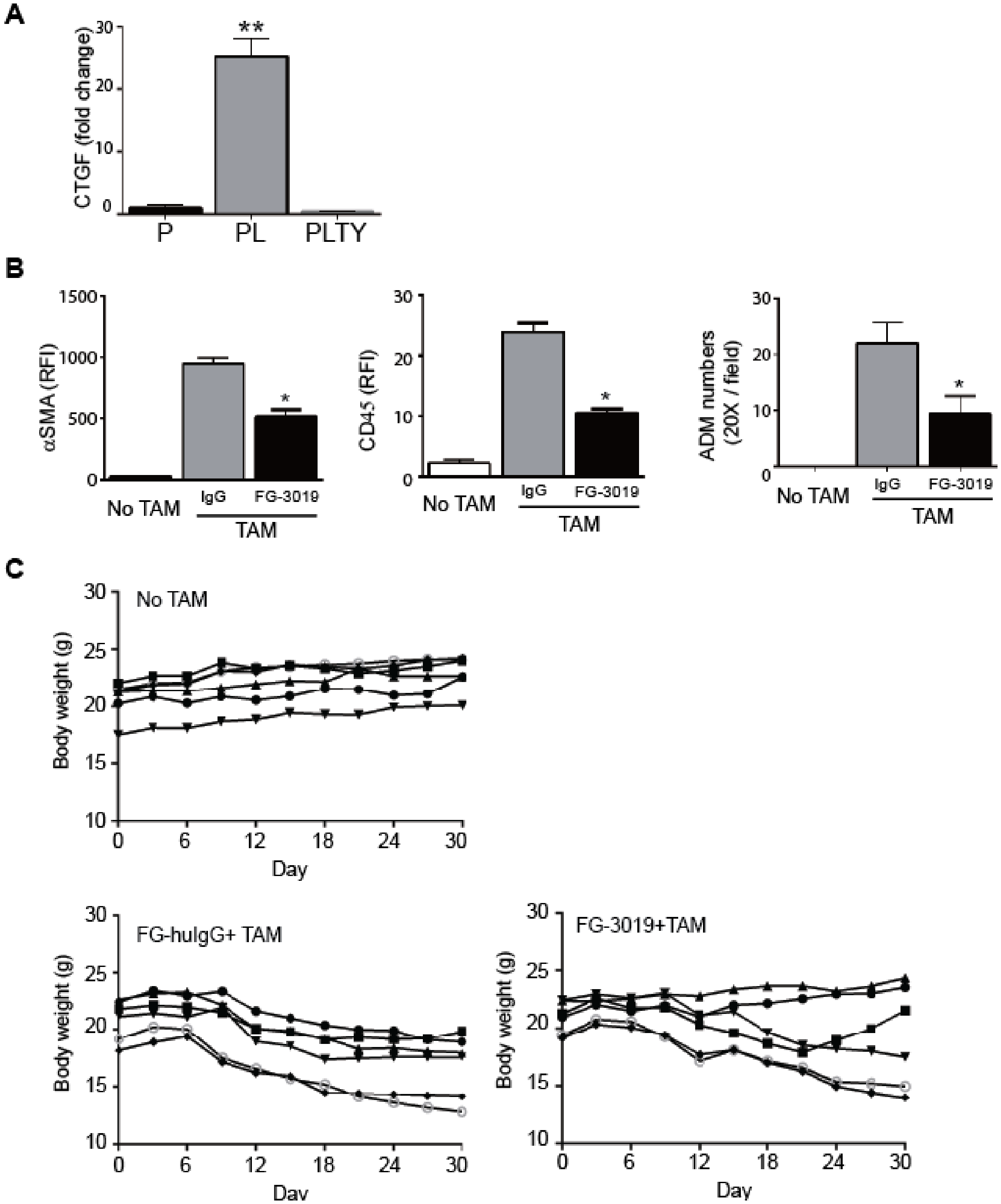

Supplementary Figure 10

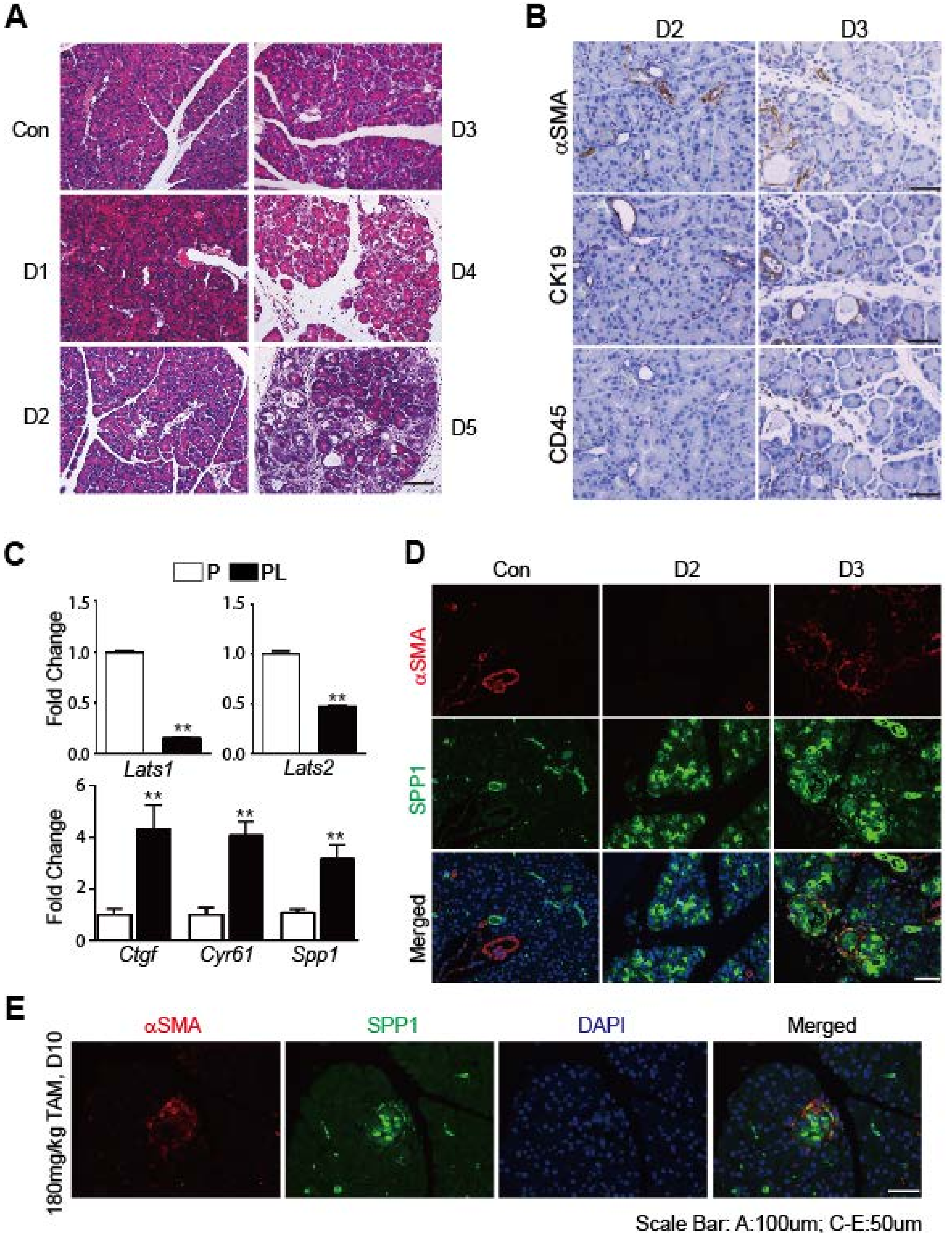

Supplementary Figure 11

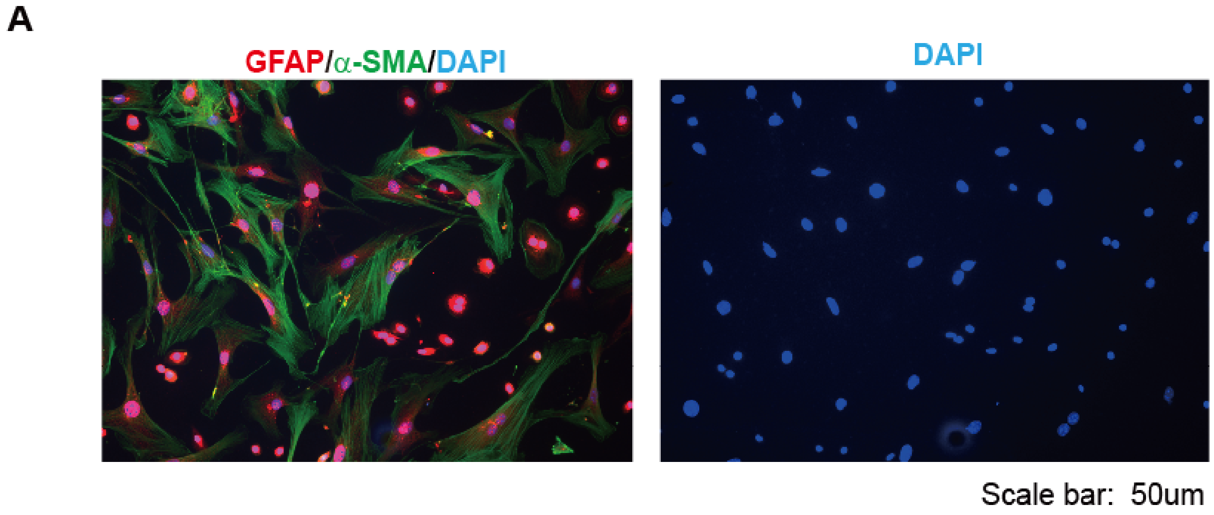

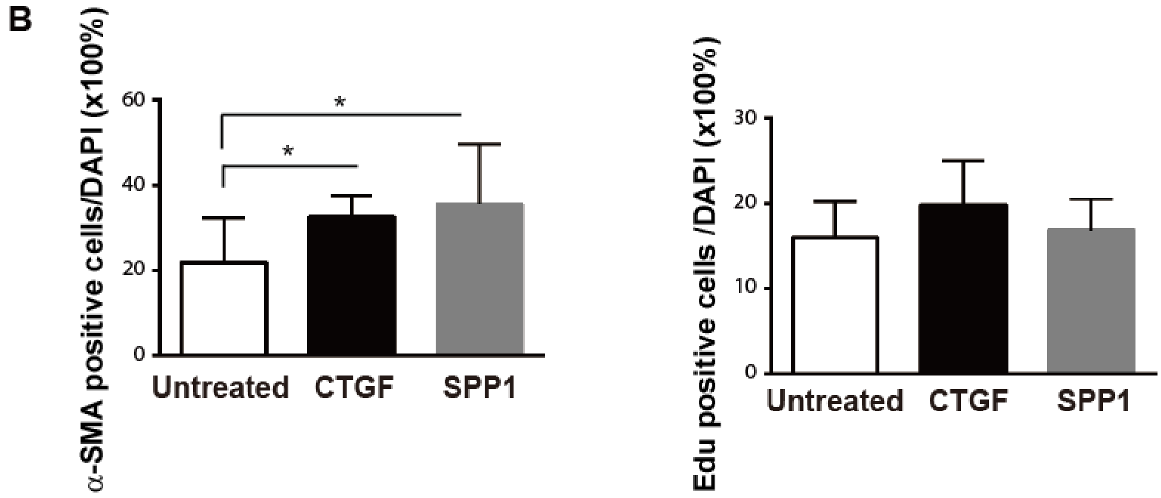

Supplementary Figure 12

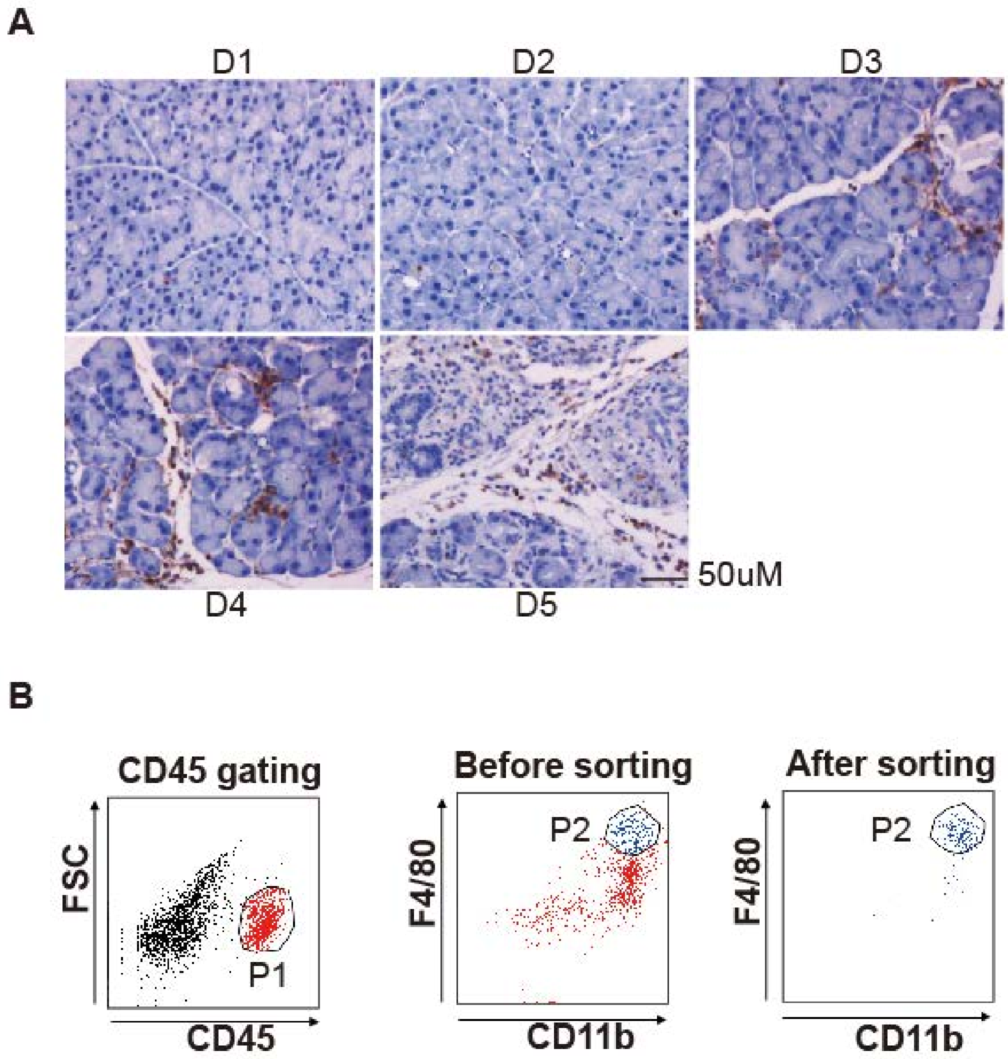
